# Repurposing covalent EGFR/HER2 inhibitors for on-target degradation of human Tribbles 2 (TRIB2) pseudokinase

**DOI:** 10.1101/305243

**Authors:** Daniel M Foulkes, Dominic P Byrne, Fiona P Bailey, Samantha Ferries, Claire E Eyers, Karen Keeshan, Safal Shrestha, Wayland Yeung, Natarajan Kannan, Carrow Wells, David H Drewry, William J Zuercher, Patrick A Eyers

## Abstract

**ONE SENTENCE SUMMARY:** A Tribbles 2 pseudokinase small molecule screen led to the identification of known EGFR/HER2 inhibitors that alter the stability of TRIB2 *in vitro* and lead to rapid on-target degradation of TRIB2 in human cancer cells.

**SHORT ABSTRACT:** Tribbles 2 (TRIB2) is a cancer-associated pseudokinase with a diverse interactome, including the AKT signaling module. Substantial evidence demonstrates that TRIB2 dysregulation is important in multiple human tumors. The non-canonical TRIB2 pseudokinase domain contains a unique cysteine rich region and interacts with a peptide motif in its own C-terminal tail. We demonstrate that TRIB2 is a target for previously described small molecule protein kinase ‘inhibitors’, which were originally designed to inhibit the catalytic domain of EGFR/HER2 tyrosine kinases. Using thermal-shift assays and drug repurposing, we classify ligands that stabilize or destabilize the TRIB2 pseudokinase domain. TRIB2 destabilizing agents, including the clinical inhibitor afatinib, lead to rapid and on-target TRIB2 protein degradation in tumor cells, eliciting tractable effects on cell signaling and survival. Our data identifies leads for further development of TRIB2-degrading drugs and highlights compound-induced TRIB2 downregulation, which might be mechanistically relevant for other catalytically-deficient (pseudo)kinases targeted by small molecules.

**FULL ABSTRACT:** A major challenge associated with biochemical and cellular analysis of pseudokinases is the lack of target-validated small molecule ligands with which to probe molecular function. Human Tribbles 2 (TRIB2) is a cancer-associated pseudokinase with a diverse interactome, which includes the canonical AKT signaling module. There is substantial evidence that human TRIB2 is a therapeutic target in both solid tumors and blood cancers. The non-canonical TRIB2 pseudokinase domain contains a unique cysteine-rich region and interacts with a peptide motif in its own C-terminal tail, which was previously shown to drive interaction with cellular E3 ubiquitin ligases. In this study we demonstrate that TRIB2 is a target for previously described small molecule protein kinase inhibitors, which were originally designed to inhibit the canonical catalytic domain of the tyrosine kinases EGFR/HER2. Using a thermal-shift assay, we discovered TRIB2 ligands within the Published Kinase Inhibitor Set (PKIS), and employed a drug repurposing approach to classify compounds that either stabilize or destabilize TRIB2 *in vitro*. Remarkably, TRIB2 destabilizing agents, including the clinical covalent drug afatinib, lead to rapid and on-target TRIB2 degradation in human cells, eliciting tractable effects on signaling and survival. Our data reveal the first drug-leads for development of TRIB2-degrading ligands, which will also be invaluable for unravelling the cellular mechanisms of TRIB2-based signaling. Our study highlights that small molecule-induced protein downregulation through drug ‘off-targets’ might be relevant for other inhibitors that serendipitously target pseudokinases.

**ABBREVIATIONS**
DSF: Differential Scanning Fluorimetry
**EGFR**: Epidermal Growth Factor Receptor
**HER2**: Human Epidermal Growth Factor Receptor 2
**MS**: Mass spectrometry
**MST**: MicroScale Thermophoresis
**PKIS**: Published Kinase Inhibitors Set
**TRIB2**: Tribbles 2
**TSA**: Thermal Stability Assay

## INTRODUCTION

The human protein kinome encodes ∼60 protein pseudokinases, which lack at-least one conventional catalytic residue, but often control rate-limiting signaling outputs within cellular networks [1]. Like canonical kinases, pseudokinases drive conformation-dependent signaling associated with normal physiology and disease [2, 3]. The human ‘pseudokinome’ includes cancer-associated signaling proteins such as HER3, JAK2 (JH2 domain) and TRIB2, which have historically received much less attention compared to their conventional, catalytically-active, counterparts even though pseudokinase domains represent rational targets for drug discovery [4]. Discovering or repurposing biologically and/or clinically-active ligands that target atypical-conformations of canonical kinases or pseudokinases, is an area of active research [2, 3, 5–9]. Moreover, the burgeoning pseudokinase field is strongly placed to benefit from the decades of research undertaken on canonical protein kinases, which has seen the approval of over 40 kinase inhibitors for human cancer and inflammatory diseases [10, 11].

The three human Tribbles (TRIB) pseudokinases are homologues of the *Drosophila melanogaster* pseudokinase termed Tribbles, which controls ovarian border cell and neuronal stem cell physiology [12, 13]. Tribbles, the human TRIB1, 2 and 3 orthologues, and the related pseudokinase STK40, all contain a catalytically-impaired pseudokinase domain. Adaptions in the pseudokinase fold, including a highly unusual αC-helix, are thought to support a competitive regulatory interaction *in cis* with a unique C-terminal tail DQLVP motif [14–16]. Through a still obscure mechanism, TRIB and STK40 function as adaptor proteins that recruit ubiquitin E3 ligases such as COP1 through direct interaction with the conserved C-tail peptide [14, 16], which is required for signaling and cellular transformation [17]. Mechanistically, Tribbles signaling outputs are controlled through the ubiquitylation and subsequent proteasomal destruction of Tribbles ‘pseudosubstrates’, such as vertebrate C/EBPα, CDC25C and Acetyl CoA carboxylase [18–20].

A longstanding goal in cancer research is drug-induced degradation of oncogenic proteins. Progress towards this objective has been transformed by the synthesis of proteolysis-targeting chimeras (PROTACs), which induce proteasome-dependent degradation of their targets. Multifunctional small molecule PROTACS often possess ligand-binding regions derived from kinase inhibitors [21, 22], and multiple classes of non-PROTAC kinase inhibitor also induce kinase target degradation, although typically at higher (micromolar) concentrations than those required for enzymatic inhibition [7]. Recent reports also disclose classes of covalent ligands that bind and disable Cys-containing small G-proteins such as mutant human RAS, permitting covalent inactivation of this previously ‘undruggable’ oncoprotein [23, 24]. Cys residues are widespread and highly conserved in kinases [25] and the conservation of Cys residues both inside and outside the catalytic domain provides kinome-wide opportunities for exploitation using chemical biology [26]. In this context, covalent targeting methodologies involving compound-accessible Cys residues in kinases [8, 27–30] and pseudokinases [3], have attracted significant attention for small molecule design, due to the potential for gains in target specificity and durability of responses, combined with tractability in experimental systems.

Tribbles pseudokinases are implicated in a huge variety of physiological signalling pathways, often in the context of protein stability, but also through regulation of key modules, such as the canonical AKT pathway [31]. TRIB2 is also implicated in the aetiology of human cancers, including leukemias, melanoma, lung and liver cancer [32]. In particular, TRIB2 is a potential drug target in subsets of acute myeloid and lymphoid leukemia (AML and ALL), which are in urgent need of targeted therapeutics to help treat untargeted or drug-resistant patient populations [33]. TRIB2 protein levels have also been linked to drug-resistance mechanisms, where an ability to modulate the pro-survival AKT signaling module underlies a central regulatory role in cell proliferation, differentiation, metabolism and apoptosis [31, 34–38].

In this paper, we report that the low-affinity TRIB2 ATP-binding site [39] is druggable with small molecules previously recognized as ATP-competitive pan-EGFR/HER2 kinase inhibitors. Biochemical analysis confirms the existence of distinct ligand-induced TRIB2 conformations and a compound screen identifies known EGFR/HER2 inhibitors that stabilize or destabilize TRIB2 *in vitro*. TRIB2 ligands include the clinical breast cancer therapeutic lapatinib/Tykerb [40] and the approved irreversible electrophilic covalent inhibitors afatinib/Giotrif [41, 42] and neratinib/Nerlynx [43, 44]. In the case of these two destabilizing agents, binding leads to uncoupling of the pseudokinase domain from its own C-terminal tail. Consistently, cellular afatinib exposure leads to rapid TRIB2 degradation driven by an interaction with the Cys-rich pseudokinase domain, which interferes with AKT signaling and decreases cell survival in a TRIB2-expressing leukemia model. The availability of target-validated ligands that act as rapid TRIB2 pseudokinase down-regulators through a direct effect on the pseudokinase, represents a new way to evaluate TRIB2 physiology and cell signaling. It might also have a broader impact on the rapidly developing pseudoenzyme field [45], where the concept of pseudokinase destabilization or elimination by targeted kinase ‘inhibitors’ [7] has a number of potentially useful applications.

## RESULTS

### Analysis of human TRIB2 using a thermal stability assay

Human TRIB2 differs from TRIB 1 and TRIB3 in the pseudokinase domain due to a Cys-rich region at the end of the β3 Lys-containing motif, which extends into the truncated αC-helix in the N-lobe (Figure 1A, top). We developed a Differential Scanning Fluorimetry (DSF) assay [46–48] to examine thermal stability of full-length (1-343) His-tagged TRIB2 protein, and compared it to full-length cAMP-dependent protein kinase (PKAc) catalytic subunit, or full-length Cys 104 Tyr TRIB2, in which Cys 104 was replaced with the Tyr residue conserved in human TRIB1 and TRIB3 (Figure 1A). Proteins were purified to homogeneity (Figure 1A, bottom) and thermal stability based on unfolding profiles were determined for each protein (reported as a T_m_ value, Figure 1B). As previously demonstrated [39], TRIB2 (T_m_ = ∼39 °C) was much less thermostable than the canonical protein kinase (PKA, T_m_ = 46.3 °C). Remarkably, single substitution of Cys 104 with Tyr induced a potent stabilization of TRIB2, with the T_m_ value increasing to ∼49 °C, comparable to that of human TRIB1 [14], suggesting an important structural role for this unique Cys residue in TRIB2 (un)folding dynamics. To confirm that recombinant TRIB2 binds to a known physiological target, we demonstrated that GST-tagged TRIB2 interacted with a marked preference for catalytically inactive (non Thr 308-phosphorylated) AKT1 *in vitro* (Figure 1C). Consistent with a functional regulatory interaction between TRIB2 and AKT in cells [49], transient overexpression of Tet-inducible FLAG-tagged TRIB2 in HeLa cells led to a marked increase in endogenous AKT phosphorylation at the hydrophobic motif (Ser 473, Figure 1D), an established marker for AKT catalytic activity and the downstream generation of a cellular anti-apoptotic signal [50].

**Figure 1.**
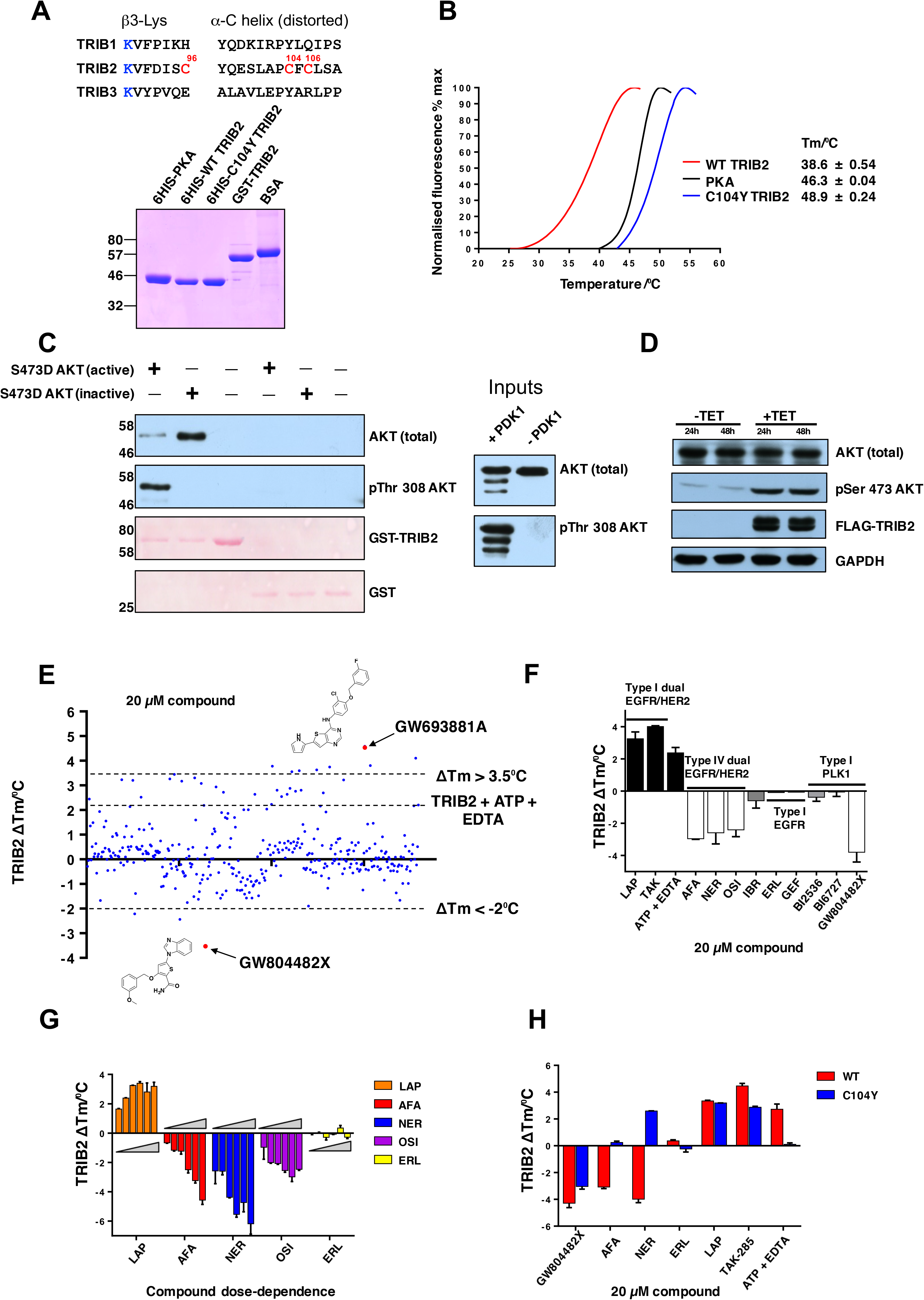
Full-length TRIB2 is a target for protein kinase inhibitors *in vitro*. **(A)** Sequence alignment of human TRIB1, TRIB2 and TRIB3, highlighting Cys-rich residues (red) in the TRIB2 pseudokinase domain. Recombinant proteins employed for *in vitro* analysis. 5 μg of the indicated purified proteins were resolved by SDS-PAGE. **(B)** Thermal denaturation profiles of recombinant proteins. A representative unfolding profile is shown. T_m_ values (±SD) were obtained from 3 separate fluorescence profiles, each point assayed in duplicate. **(C)** The ability of GST-TRIB2 to interact with active (PDK1-phosphorylated) or inactive (non PDK1-phosphorylated) Ser 473 Asp AKT1 was assessed by GSH-sepharose pull-down followed by immunoblotting. PDK1 phosphorylated Ser 473 Asp AKT is also phosphorylated on Thr 308 (right panel), but binds very weakly to TRIB2. **(D)** Transient transfection of Tet-inducible FLAG-TRIB2 leads to increased AKT phosphorylation on Ser 473. **(E)** TRIB2 DSF screen using PKIS. 5 μM His-TRIB2 was employed for all DSF analysis. ΔT_m_ values were calculated for each compound. Scattergraph of data highlights a wide variety of compounds that either stabilize or destabilize TRIB2 *in vitro*. Cut off values of >3.5 °C and < 2°C were used to designate ‘hits’. **(F)** Comparative DSF analysis of clinical and preclinical kinase inhibitors as potential TRIB2 ligands. LAP=lapatinib, TAK=TAK-285, AFA=afatinib, NER=neratinib, OSI=osimertinib, IBR=ibrutinib, ERL=erlotinib, GEF=gefitinib. **(G)** Dose-dependent analysis of thermal shifts induced by clinical TRIB2 ligands. Compounds were tested at 5, 10, 20, 40, 80 and 160 μM. **(H)** Profiling of TRIB2 and C104Y with selected inhibitors by DSF.

### A DSF screen for TRIB2 ligands using a kinase inhibitor library

The ability of full-length recombinant human TRIB2 to bind to ATP in the presence of EDTA [39, 47] confirms that a vestigial nucleotide-binding site is present within the pseudokinase domain. Moreover, our previous work established that an analogue-sensitive (Phe 190 Gly) TRIB2 variant could be stabilized by ‘bumped’ pyrimidine analogues *in vitro* [39]. In order to discover drug-like ligands for WT full-length TRIB2, we screened the Published Kinase Inhibitor Set (PKIS), a collection of high-quality class-annotated kinase inhibitors [51]. We enforced cut off values of ∼ΔT_m_ = <-2 °C and >+3.5 °C (therefore eliminating ∼97% of the library) to define ‘hit’ compounds that possessed the ability to either destabilize or stabilize TRIB2 respectively in a thermal stability assay (TSA) at a 1:4 TRIB2:compound molar ratio (Figure 1E and Supplementary Table 1). The top ‘stabilizing’ compound identified was GW693881A, a dual EGFR/HER2 thienopyrimdine inhibitor exhibiting a ΔTm of +4.7 °C. The top ‘destabilizing’ compound was GW804482X, a thiophene polo-like kinase (PLK) inhibitor that induced a ΔT_m_ of – 3.4 °C (Figure 1E, red symbols). Interestingly, most of the top stabilizing and destabilizing compounds belong to well-known ATP-competitive pyrimidine or quinazoline EGFR/HER2 chemotypes (Supplementary Figure 1) [52–54], suggesting broad structural cross-reactivity between TRIB2 and the ligand-responsive EGFR/HER2 conformation. To build upon these findings, we screened a larger panel of known dual EGFR/HER2 inhibitors (Supplementary Figure 2), and established that the clinical type I EGFR/HER2 inhibitors TAK-285 and lapatinib also stabilized TRIB2 *in vitro* (Figure 1F). Interestingly, the ATP-competitive covalent EGFR/HER2 inhibitors afatinib, neratinib and osimertinib (but not the unrelated covalent Bruton’s Tyrosine Kinase (BTK) inhibitor ibrutinib or the Type I EGFR-specific inhibitors erlotinib or gefitinib) destabilized TRIB2, similar to the PLK inhibitor GW804482X (Figure 1F) and the dual EGFR/HER2 inhibitor GW569530A (Supplementary Figure 1). Compound effects were caused through TRIB2 targeting in the TSA, because no shift was elicited when PKAc was compared in a side-by-side counter-screen, using dasatinib as a positive control (Supplementary Figure 3). Interestingly, the preclinical PLK inhibitors BI2536 and BI6727 (volasertib) had no discernible effect on TRIB2 stability in this assay, in contrast to GW804482X, a PLK inhibitor originally isolated in the screen (Figure 1F).

Afatinib, neratinib and osimertinib are all covalent (type IV) inhibitors of canonical EGFR/HER2 tyrosine kinase targets, interacting irreversibly with a conserved Cys residue in the ATP-binding site [55, 56]. As show in Figure 1G, the stabilization of TRIB2 by lapatinib, and the destabilization of TRIB2 by all three covalent EGFR/HER2 inhibitors, occurred in a dose-dependent manner. Interestingly, a Cys 104 Tyr TRIB2 mutant was no longer destabilized by either afatinib or neratinib, but remained ‘sensitive’ to lapatinib TAK-285 and GW804482X based on thermal-shifting (Figure 1H). Importantly, none of the latter compounds contain an electrophilic ‘warhead’ required for covalent interaction (Supplementary Figure 2). Cys 104 Tyr TRIB2 was also insensitive to thermal shift in the presence of ATP and EDTA (Figure 1H and Supplementary Figure 4). These data suggest that Cys 104 is both a key determinant for ATP binding and TRIB2 interaction with covalent kinase inhibitors, which induce TRIB2 destabilization *in vitro*.

### Mechanistic analysis of TRIB2 structural stability and TRIB2 ligand binding

Structural analysis of TRIB1 by X-ray crystallography and small angle X-ray scattering led to the proposal of an *in cis* self-assembly model, whereby the unique C-terminal tail region, which contains the conserved ‘DQLVP’ motif, binds directly to the pseudokinase domain adjacent to the short αC-helix of TRIB1 [14, 15]. We exploited DSF to investigate whether this mechanism is also relevant in TRIB2 by generating a series of truncated proteins. These lacked either the N-terminal extension, which was predicted to be disordered by both I-TASSER [57] and VSL2 [58], the C-terminal tail, or both N and C-terminal domains. We also generated a triple point mutant in which the ‘DQLVP’ tail motif, which is required for TRIB1 and TRIB2 cellular transformation *in vivo* [17], was mutated to ‘AQLAA’ (Figure 2A). Full-length (1-343) TRIB2 and TRIB2 lacking the N-terminal domain (TRIB2 54-343) both exhibited similar Tm values of 39-40 °C (Figure 2B). In contrast, deletion of the C-tail (TRIB2 1-318) markedly changed TRIB2 stability, with T_m_ values falling to ∼37°C, diagnostic of a destabilized TRIB2 conformation (Figure 2B). Mutation of DQLVP to AQLAA further destabilized TRIB2, inducing a T_m_ value of ∼36 °C (Figure 2B). Using this panel of recombinant TRIB2 proteins, we measured the relative effects of kinase inhibitors on TRIB2 stability. Consistent with a lack of effect on compound interactions, deletion of the TRIB2 N-terminal region had no effect on ΔT_m_ values induced by any of the compounds. Removal of the C-tail region (54-318 and 1-318 mutants) completely abolished afatinib and neratinib-induced TRIB2 destabilization, but had a negligible effect on GW804482X binding (Figure 2C). Consistently, destabilization by afatinib was also completely abolished in the AQLAA triple mutant, whereas neratinib effects were reduced by >50%. Notably, neither the destabilizing effect of GW804482X, nor the stabilizing effects of lapatinib or TAK-285 differed between any of the TRIB2 proteins evaluated. These results suggest a destabilizing mechanism induced by covalent EGFR/HER2 ligands through displacement of the TRIB2 C-tail, a unique feature of Tribbles pseudokinases [12, 14].

**Figure 2.**
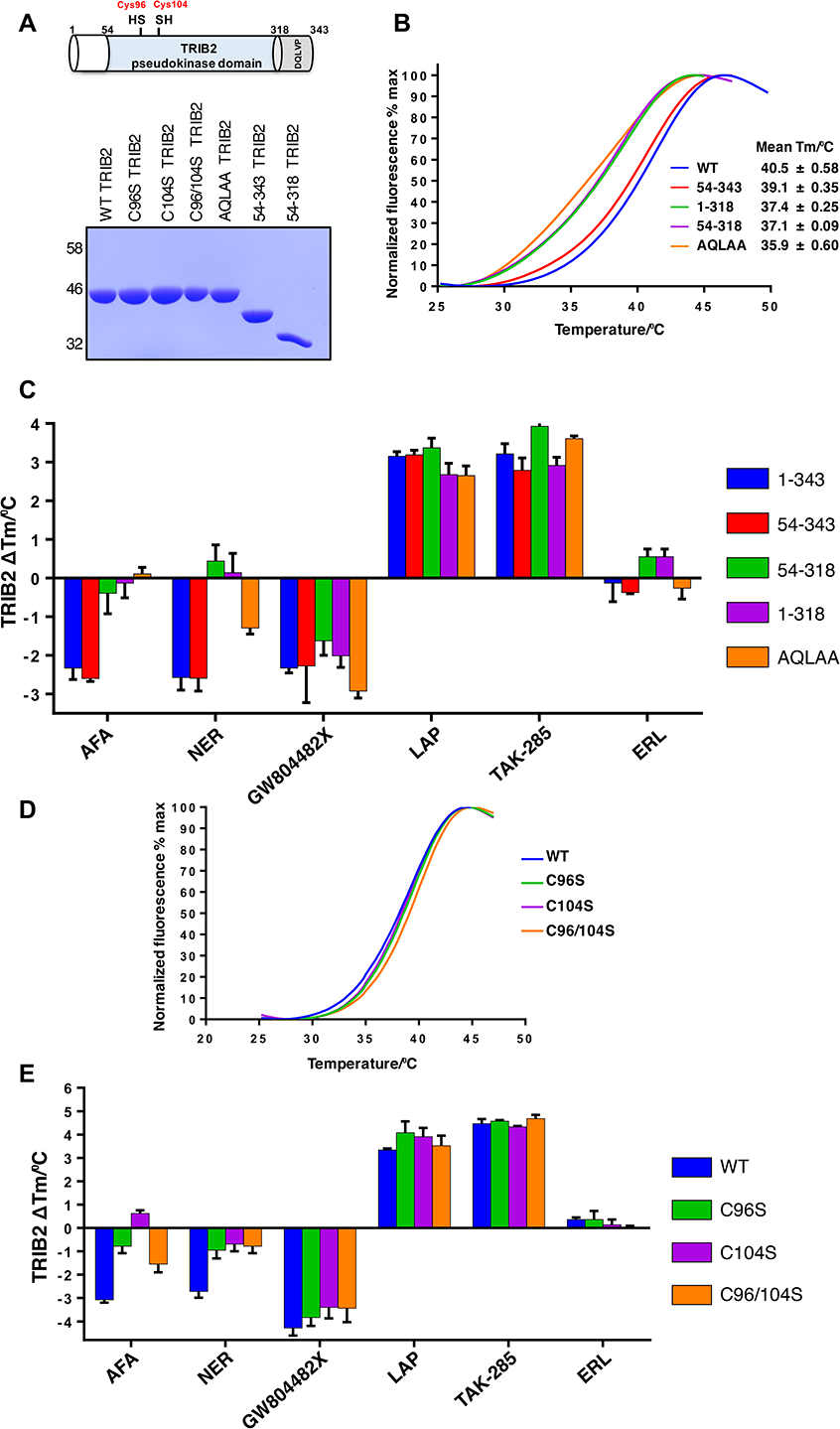
TRIB2 thermal stability is modulated through Cys binding to covalent inhibitors. **(A)** Schematic cartoon of TRIB2 with domain boundaries numbered and cysteine residues highlighted (red). SDS-PAGE of 5 μg of recombinant TRIB2 proteins. **(B)** Thermal denaturation profiles of 5 μM WT-TRIB2 (amino acids 1-343), three truncated variants and an AQLAA triple-point mutant. Representative curves for each protein and average Tm values (±SD) are shown. **(C)** Thermal shift analysis of TRIB2 deletion and AQLAA proteins measured in the presence of a panel of compounds (20 μM). The change in Tm value (ΔT_m_) is reported from duplicate experiments, each performed in triplicate and the chemical structures of each compound are shown for comparison. AFA=afatinib, NER=neratinib, LAP= lapatinib, ERL=erlotinib. **(D)** Thermal denaturation profiles for purified TRIB2 and Cys 96 Ser, Cys 104 Ser and Cys 96/104 Ser proteins. **(E)** Thermal shift analysis of TRIB2 Cys-mutated proteins measured in the presence of a panel of compounds (20 μM). The change in Tm value (ΔT_m_) is reported from triplicate experiments.

To evaluate whether the EGFR/HER2 ligand afatinib targeted the unique Cys residues in the TRIB2 pseudokinase domain (Figure 1A), we performed mass spectrometry (MS) analysis to evaluate protein modification. As shown in Supplementary Figure 5A, incubation of TRIB2 with a 5-fold molar excess of afatinib led to covalent afatinib interaction with Cys 96. A doubly charged chymotryptic product ion representing the TRIB2-derived DISC^96^Y:afatinib peptide adduct at *m/z* 543.2 (Supplementary Fig. 5A) and the isotopic ratios of the ^35^Cl or ^37^Cl-containing peptide ions unequivocally confirmed Cys 96 as a site of TRIB2 binding (Supplementary Figure 5B). Having confirmed an intact mass for recombinant full-length TRIB2 of 43,587.09 Da, very similar to the predicted mass of 43,587.22 Da (Supplementary Figure 5C), we were also able to ascertain that pre-incubation with afatinib generated covalent adducts containing predominantly either 1 or 2 molecules of afatinib. There was also some evidence for tri and tetra-modified TRIB2 adducts (Supplementary Figure 5D). We next examined afatinib interaction with TRIB2 by Microscale Thermophoresis (MST), a biophysical technique for quantification of reversible biomolecular interactions [59]. This confirmed an interaction between afatinib and fluorescent NTA-coupled His-TRIB2, which could be fitted to a reversible binding event with a K*d* value of ∼16 μM (Supplementary Figure 6). Moreover, it suggested initial (reversible) binding of the ATP-competitive afatinib ligand [60, 61] prior to subsequent formation of a covalent TRIB2 adduct(s) (Supplementary Figure 5). Finally, we evaluated the potential interaction of afatinib and neratinib with a modelled TRIB2 pseudokinase domain [12] using AutoDock Vina [62]. Blind docking of the covalent inhibitors employing the entire TRIB2 homology model revealed a putative binding pocket formed by residues in the vestigial C-helix, including Cys 96, the P-loop and the activation loop. We assessed potential binding modes for afatinib and neratinib using a covalent docking approach [63], in which the compounds were flexibly docked to the identified pocket by covalently linking the electrophilic warhead to the sulfhydryl Cys. Covalent docking revealed structurally and energetically-feasible binding modes for both afatnib and neratinib bound to Cys 96. The most commonly sampled poses are shown in Supplementary Figure 7.

To validate the functional importance of Cys residues for afatinib binding, we examined the interaction of this compound with TRIB2 in which two Cys amino acids were mutated to non-thiol containing Ser residues. Individual or combined mutation of Cys 96 and Cys 104 to a Ser residue had no effect on the thermal profile (T_m_) of the purified TRIB2 proteins (Figure 2D), in contrast to the highly stabilizing effect of a Tyr at position 104 (Figure 1B and Supplementary Figure 4). However, individual or joint mutation of Cys 96 and Cys 104 to Ser severely blunted the destabilizing effect of afatinib and neratinib on TRIB2, in contrast to the non-covalent TRIB2 ligand GW804482X or the EGFR/HER2 stabilizing compounds lapatinib and TAK-285 (Figure 2E). Taken together, these findings confirm that several EGFR/HER2 compounds bind to TRIB2 through Cys96 and/or Cys104.

### Evaluation of TRIB2 ligands in human cells

To evaluate the effects of various kinase inhibitors on TRIB2 in living cells, we generated a polyclonal TRIB2 antibody and an isogenic stable HeLa cell line expressing Tet-inducible FLAG-tagged human TRIB2. Polyclonal TRIB2 antibodies were equally efficient at recognising recombinant His-FLAG-TRIB2 and cellular FLAG-TRIB2 as a commercial monoclonal FLAG antibody (Supplementary Figure 8A). We also constructed a stable HeLa cell line expressing low levels of inducible FLAG-TRIB2 (Supplementary Figure 8B). We estimated that ∼1 ng of TRIB2 was expressed in 40 μg of whole cell lysate in the presence of Tet, by comparing known amounts of recombinant His-FLAG-TRIB2 protein (Supplementary Figure 8B).

To quantify effects of TRIB2 expression on canonical signaling in model cells, we first evaluated AKT phosphorylation. As shown in Figure 3A, AKT became phosphorylated in stable HeLa cells after serum stimulation in the absence of Tet. Tet-induced TRIB2 expression led to the appearance of a TRIB2 doublet (phosphorylation confirmed in the upper band by lambda phosphatase treatment, Supplementary Figure 9), and a marked increase in the extent and duration of AKT phosphorylation at Ser 473. Using this HeLa cell model, we next evaluated the effects of small molecule inhibitors on TRIB2 signaling, by comparing a panel of *in vitro* TRIB2 destabilizing ligands discovered by DSF *in vitro* (afatinib, neratinib and osimertinib) with a series of control compounds. Brief (4h) exposure of TRIB2-expressing cells to afatinib led to a specific decrease in TRIB2 protein expression. All of the compounds evaluated are EGFR or EGFR/HER2 signaling pathway inhibitors, and we confirmed they all blocked ERK phosphorylation at these concentrations (Figure 3B); any unique effects of drugs in this panel are therefore likely to be ‘off-target’ to EGFR/HER2. We next incubated cells for increasing lengths of time with afatinib. A rapid, time-dependent, elimination of TRIB2 protein was evident when cells were exposed to this drug, in contrast to DMSO controls in which TRIB2 protein accumulated over time (Figure 3C), as expected [18]. To evaluate intracellular interaction between TRIB2 and kinase inhibitors, we exposed HeLa cells expressing Tet-inducible FLAG-TRIB2 to each compound, and quantified TRIB2 thermal stability in the cell extracts using a cellular TSA (Supplementary Figure 10) [64]. Consistently, FLAG-TRIB2 was destabilized more rapidly than DMSO and erlotinib-treated controls in the presence of afatinib, becoming undetectable in extracts heated to 45 °C (Supplementary Figures 10A and 10B). This was distinct from lapatinib, which partially stabilized exogenous TRIB2 in cell extracts, consistent with *in vitro* analysis (Figure 1F).

**Figure 3:**
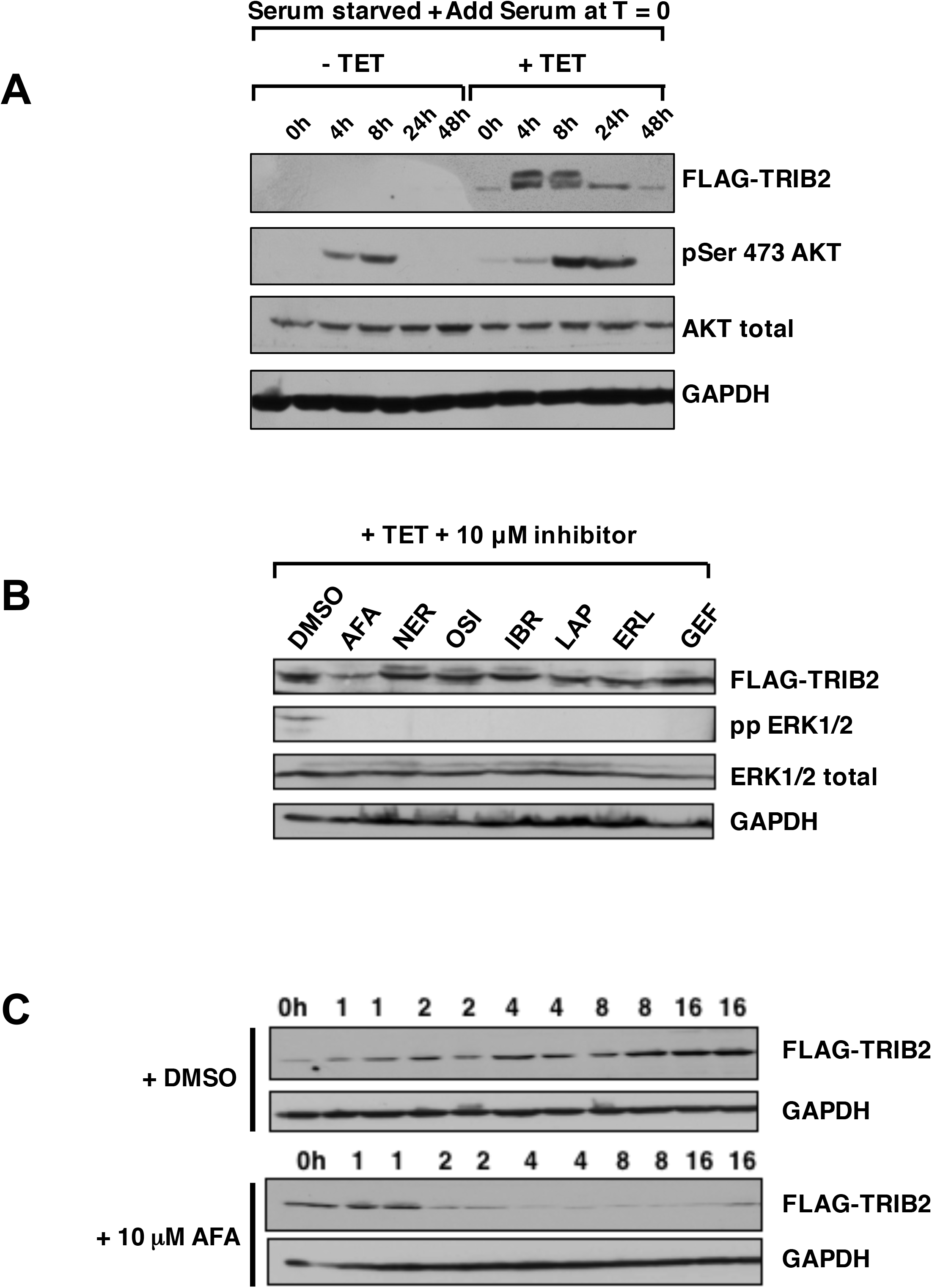
Afatinib promotes rapid degradation of FLAG-TRIB2 in an inducible HeLa cell model. **(A)** Uninduced (-Tet) or Tetracycline-induced (+Tet) HeLa cells containing a stably integrated FLAG-TRIB2 transgene were serum starved for 12 h prior to the addition of serum, and lysed at the indicated times. Whole cell extracts were blotted with FLAG antibody to detect FLAG-TRIB2, pSer 473 AKT. Total AKT and total GAPDH served as loading controls. **(B)** A selection of clinically approved kinase inhibitors, including dual EGFR/HER2 and EGFR-specific compounds, were added to Tet-induced cells at a final concentration of 10 μM. Stable cells were induced to express FLAG-tagged TRIB2 with tetracycline for 16h prior to inhibitor treatment for 4 hours. AFA=afatinib, NER=neratinib, OSI=osimertinib, IBR=ibrutinib, LAP=lapatinib, ERL=erlotinib, GEF=gefitinib. Whole cell extracts were immunoblotted with FLAG, pERK, ERK or GAPDH antibodies. **(C)** Stable HeLa cells were incubated with Tet for 16h, and then incubated with DMSO or 10 μM afatinib (AFA) prior to lysis at the indicated time. Whole cell extracts were immunoblotted with FLAG or GAPDH antibodies.

Afatinib interacts with TRIB2 through a biochemical Cys-based mechanism, so we generated isogenic Tet-inducible Cys 96 Ser or Cys 96/104 Ser TRIB2 mutant stable cell lines and evaluated the effects of afatinib on exogenous TRIB2 stability. As shown in Figure 4A, afatinib (but not lapatinib or TAK-285) induced dose-dependent loss of TRIB2 in WT-TRIB2 cells, which was partially prevented by mutation of Cys 96 to Ser, and completely abolished in double Cys-expressing cells (quantified in Figure 4A, right panels), demonstrating unequivocally that afatinib binds to TRIB2 in these cells. Furthermore, afatinib (but not TAK-285 or erlotinib) treated WT-TRB2 cells exhibited a marked decrease in AKT Ser 473 phosphorylation, and this effect was abrogated in double Cys-expressing mutant cells (Figure 4B). Importantly, afatinib, TAK-285 and erlotinib all blocked ERK phosphorylation in WT and C96/104S-TRIB2 isogenic lines, consistent with ‘on-target’ inhibition of their common target EGFR (Figure 4B).

**Figure 4.**
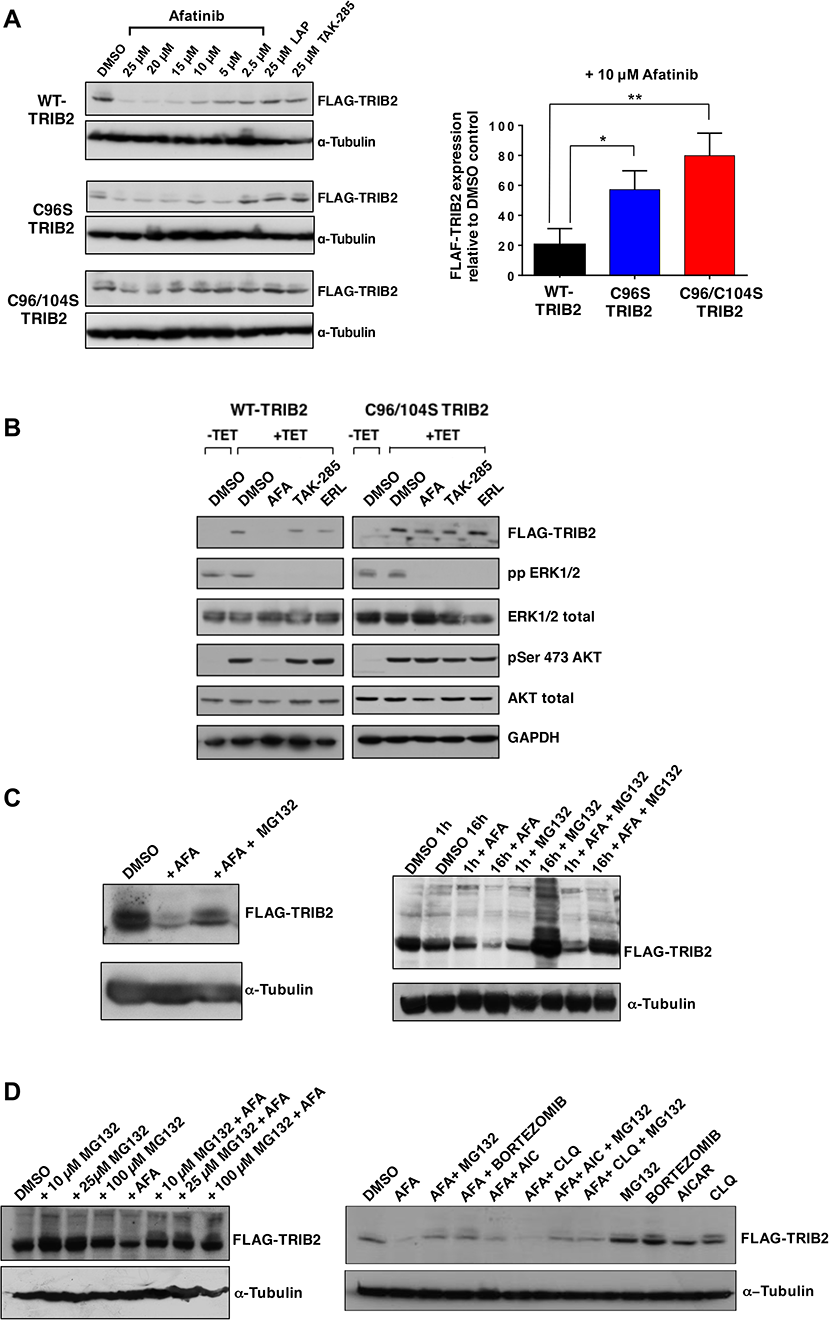
On-target degradation of TRIB2 by afatinib: Cys 96/104 Ser TRIB2 is resistant to degradation. **(A)** The indicated concentration of afatinib, lapatinib or TAK-285 (10 μM) was incubated for 4 h with isogenic stable HeLa cells expressing FLAG-tagged WT-TRIB2, Cys 96 Ser or Cys 96/104 Ser TRIB2 (induced by Tet exposure for 16h). After lysis, whole cell extracts were immunblotted with the indicated antibodies. FLAG-TRIB2 levels were quantified after exposure to 10 μM afatinib relative to DMSO controls using ImageJ densitometry software, with a representative experiment shown (significance assessed using data from three independent experiments and students t-test p=0.019 and p=0.0066 relative to FLAG-tagged WT-TRIB2). **(B)** WT and Cys 96/104 Ser stable HeLa cell lines were subjected to a serum block-and-release protocol. Subsequently, the indicated compounds (10 μM) were added for 4 h prior to cell lysis and immunoblotting with the indicated antibodies. AFA=afatinib, ERL=erlotinib. **(C)** FLAG-tagged TRIB2 expressing HeLa cells were incubated with (0.1% v/v) DMSO or 10 μM afatinib, in the presence or absence of 10 μM MG132 for 4 hours (left) or at the indicated time points (right) prior to lysis and processing for immunoblotting. **(D)** FLAG-TRIB2 expressing stable cells were incubated with the indicated concentration of MG132 in the presence or absence of 10 μM afatinib for 4 h prior to cell lysis and immunoblotting (left) or for 1h with 10 μM bortezomib (BOR), 1 mM AICAR (AIC) or 50 μM Chloroquine (CLQ) prior to the addition of afatinib (AFA, 10 μM) for an additional 4 h, prior to lysis and immunoblotting with the indicated antibodies.

To investigate the mechanism of TRIB2 destabilization by afatinib, we added the drug to WT-TRIB2 expressing HeLa cells in the presence of the proteasome inhibitor MG132, which partially rescued TRIB2 degradation after both rapid and prolonged exposure to afatinib (Figure 4C). This finding is consistent with previously reported proteasome-dependent TRIB2 turnover [65]. We further evaluated this mechanism using a range of MG132 concentrations and the clinical proteasome inhibitor bortezomib (Figure 4D) [20]; under both conditions, afatinib-mediated destabilization of TRIB2 was decreased. This effect was in contrast to afatinib-induced TRIB2 destabilization, which was still observed in the presence of non-specific inhibitors of autophagy (AICAR) and lysosomal degradation (chloroquine) (Figure 4D). Our observations are in line with published findings, which suggest that TRIB2 fate is dependent upon turnover by the ubiquitin proteasome system (UPS) in human cancer cells [18, 66]. Of further interest, the (non-covalent) TRIB2-destabilizing ligand GW804482X isolated from our screen (Figure 1), did not induce TRIB2 degradation in cells after a 4-hour incubation period, in contrast to proteasome-dependent degradation induced by the same concentration of afatinib (Supplementary Figure 11).

High levels of TRIB2 expression drive AML *in vivo* by inhibiting myeloid differentiation and promoting proliferation [17, 34]. However, endogenous TRIB2 expression has only previously been analysed in a few cell types, in large part due to a lack of reliable TRIB2 reagents. Using our TRIB2 antibody, we evaluated endogenous expression of TRIB2 in the clinically-relevant U937 AML cell model (Figure 5A), and established that acute (4 h) exposure to afatinib, but not lapatinib or erlotinib, decreased TRIB2 protein levels in a dose-dependent manner (Figure 5A). Consistent with their ability to inhibit EGFR signaling, all three compounds completely blocked ERK phosphorylation. We next established dose-dependent effects on both TRIB2 expression and AKT Ser 473 phosphorylation in afatinib-treated U937 cells (Figure 5B). Importantly, these effects were tightly correlated with apoptotic induction of caspase 3 cleavage, but only at afatinib concentrations that also induced TRIB2 degradation and concomitant loss of AKT Ser 473 phosphorylation (Figure 5B). To determine the impact of afatinib treatment on U937 cell viability we quantified cellular cytotoxicity after 72 h exposure (Figure 5C). Afatinib (and neratinib) reduced cell viability with sub-micromolar IC_50_ values, whereas EGFR inhibitors erlotinib and gefitinib and the dual EGFR/HER2 inhibitor TAK-285, were 10-20 fold less effective when compared side-by-side (Figure 5C). As these compounds did not induce TRIB2 destabilization, AKT activation or caspase 3 activation, our data suggest that the cellular TRIB2-destabilizing ligands afatinib and neratinib possess an enhanced ability to kill AML-derived cells [49].

**Figure 5.**
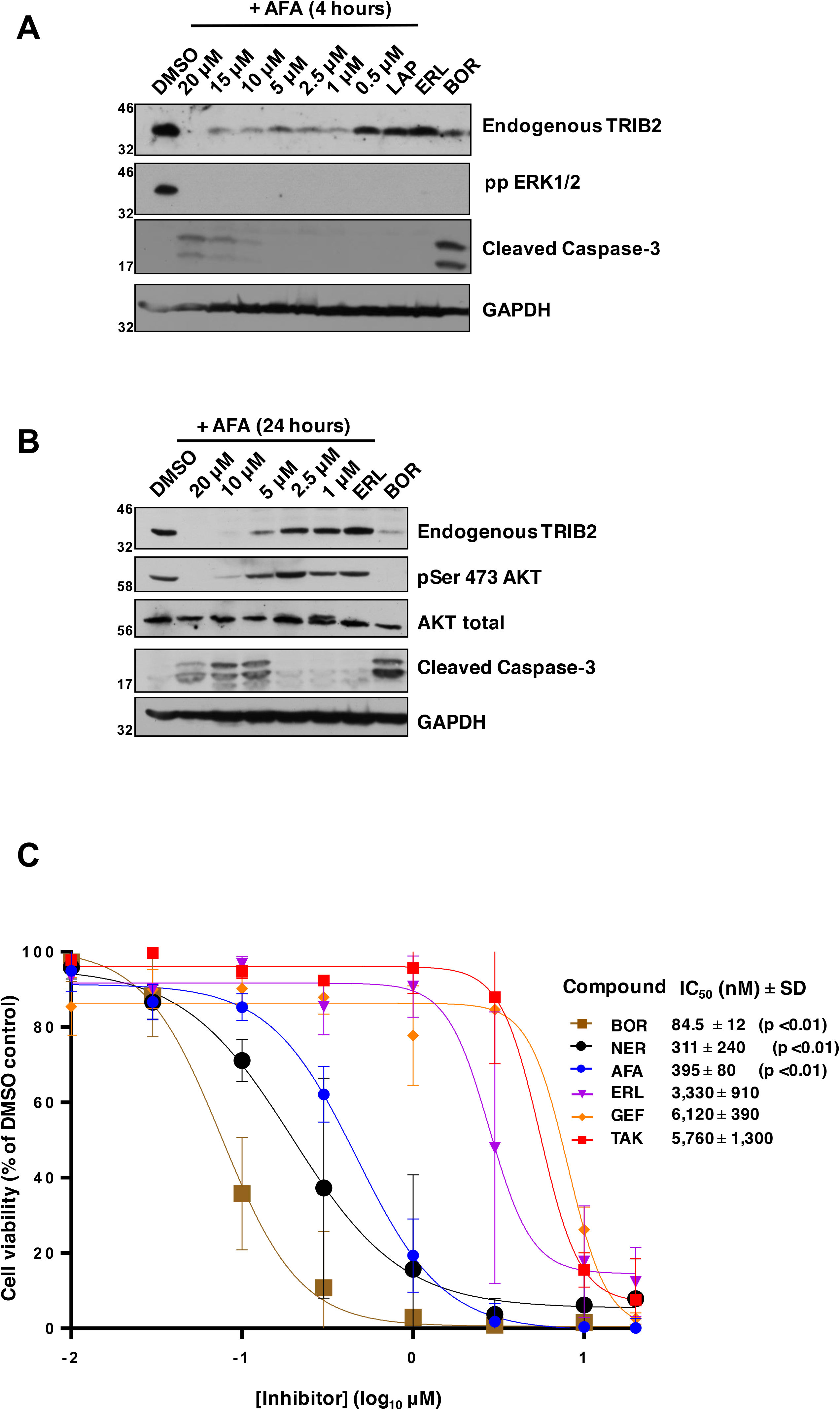
Afatinib rapidly destabilizes endogenous TRIB2 and specifically induces caspase-3 cleavage and U937 cytotoxicity. **(A)** Endogenous TRIB2 is destabilized in human U937 cells in a dose-dependent manner after a 4 h exposure to afatinib (AFA). Cells were incubated with either 0.1% (v/v) DMSO, the indicated concentrations of afatinib, or 10 μM lapatinib (LAP), 10 μM erlotinib (ERL) or 10 μM of the proteasome inhibitor bortezomib (BOR) for 4 h prior to lysis and immunoblotting of endogenous TRIB2, pERK or cleaved caspase 3. GAPDH served as a loading control. **(B)** Endogenous TRIB2 is destabilized after exposure to afatinib for 24h, concomitant with reduced AKT phosphorylation at Ser 473. After cell collection and lysis, whole cell extracts were immunoblotted with the indicated antibodies. AKT and GAPDH served as loading controls and ERL and BOR were used at 10 μM. **(C)** The cytotoxicity of a panel of TRIB2-destabilsing ligands (afatinib, AFA and neratinib, NER) were compared to EGFR inhibitors or the TRIB2 stabilizer TAK-285 (TAK). MTT assays were performed after 72 h compound exposure, with bortezomib (BOR) employed as a positive control. IC_50_ values (nM ± SD) are derived from 3 independent experiments performed in triplicate. Statistical analysis confirmed a significantly-different (∼20-fold increase) in cytotoxicity for AFA exposure when compared together with TAK-285, ERL and GEF (p=0.0006) and also significantly different cytotoxic effects of BOR, AFA and NER when compared to the EGFR inhibitor erlotinib (p=<0.01).

## DISCUSSION

### A ligand-targetable regulatory mechanism in human TRIB2

It is believed that rate-limiting signaling outputs are often controlled by non-enzymatic components, exemplified by mechanistically-related families of pseudoenzymes such as pseudokinases [45]. In the present study, we sought to target cancer-associated TRIB2 with a small molecule; initially the intention was to establish the validity of this approach for this class of pseudokinase, but subsequently we were also able to evaluate the effects of TRIB2 compound engagement in intact human cells. To discover TRIB2 ligands, we exploited an unbiased TRIB2 thermal shift assay [39, 47], and identified multiple chemical classes of dual EGFR/HER2 (but not monovalent EGFR) inhibitors. One group of (ATP-competitive) kinase inhibitor induced stereotypical (positive) thermal shifts in TRIB2, consistent with binding in the atypical TRIB2 nucleotide-binding site [47]. A second group of chemical ligands induced TRIB2 destabilization, which although initially unexpected, could be explained mechanistically by a compound-induced effect in which the TRIB2 pseudokinase domain became uncoupled from its own C-tail region. A similar mechanism has been proposed to explain thermally-distinct conformers that exist in TRIB1, which shares ∼70% identity with TRIB2 in the pseudokinase domain and a very similar C-tail motif [15]. Deletion of the TRIB1 C-tail also leads to the destabilization of the TRIB1 pseudokinase domain compared to full-length TRIB1 [14], similar to destabilizing effects observed for TRIB2 1-318 and 54-318 deletion mutants and an AQLAA mutant. These data allow us to propose that pseudokinase-domain docking to the C-terminal tail generates a thermally-stable conformation in both TRIB1 and TRIB2, and in the case of TRIB2, that this dynamic interaction can be targeted with ATP-competitive covalent kinase inhibitors, which induce a structural pseudokinase conformation associated with decreased stability *in vitro*.

### How do covalent EGFR/HER2 inhibitors target TRIB2?

We focused our studies on electrophilic type IV kinase inhibitors, since they were likely to bind to TRIB2 *via* a covalent mechanism, which is amenable to comparative chemical and mutational analysis [30, 67]. Using a combination of techniques, we confirmed that at-least two highly conserved TRIB2 Cys residues are targets of the ATP-dependent tyrosine kinase inhibitor afatinib [68] *in vitro*. Both TRIB2 Cys-residues are uniquely located between the ATP-binding β3-Lys residue (Lys90 in TRIB2) and the very short αC-helix. In most canonical kinases, including EGFR family members, positioning of this helix is critical for catalysis and switching between inactive and active enzyme conformations, both of which can be targeted with small molecules [69]. Dynamic helix positioning is known to be generally important for interactions with kinase inhibitors, some of which recognise ‘inactive’ conformations in kinases in which the αC helix adapts to permit compound binding, as established for HER2 [70]. The conformation of the flexible C-helix relative to the ATP binding P-loop also creates an allosteric binding pocket in canonical kinases such as ERK1/2, which can be targeted with selective non ATP-competitive inhibitors such as SCH772984 that possess slow off-rate kinetics [71]. In TRIB2, a unique disposition of Cys residues (Figure 1A) makes it vulnerable to destabilization by the EGFR/HER2 covalent inhibitors afatinib, neratininb and osimertinib *in vitro*, which all undergo Michael addition to a conserved Cys residue in the roof of the ATP-binding site of EGFR-related tyrosine kinases (Cys 797 in EGFR, Cys805 in HER2 and Cys803 in HER4) [61]. Interestingly, afatinib also demonstrates low, but detectable, affinity for the ATP-binding pseudokinase domain of HER3 [72, 73], which possesses a Ser residue (Ser 794) at the equivalent residue to Cys 797 of EGFR, and assumes a pseudo ‘active’ catalytic conformation, despite possessing exceptionally low kinase activity [73]. In the absence of a high-resolution TRIB2 pseudokinase crystal structure, we do not yet know the conformation(s) relevant for small molecule interaction, although the TRIB2 N-lobe has both shared and distinct features (notably TRIB2 Cys 104) compared to TRIB1 [14]. This is reflected biochemically by marked differences in the ability of TRIB2 to bind to ATP, which likely predisposes it to targeting by EGFR/HER2 kinase inhibitors (Supplementary Figure 7). Interestingly, none of the EGFR family members possess a Cys residue in the C-helical region conserved in TRIB2, and the overall identity between HER2 and TRIB2 in the (pseudo)kinase domain is very low indeed (∼22%). We therefore believe that our discovery of TRIB2 binding to covalent inhibitors such as afatinib owes as much to the unique availability of reactive Cys residues adjacent to an allosteric pocket in the TRIB2 pseudokinase domain as to the (presumably) lower affinity for the vestigial ATP-binding site. Our work builds on previous studies in which pseudokinase domains have been targeted with small molecule kinase ‘inhibitors’ [3]. Examples include the ligand TX1-85-1 [8], which binds covalently to Cys721 (inducing HER3 degradation in cells) and recently described non-covalent JAK2 JH2 (pseudokinase) domain ligands, confirmed through structural analysis [74, 75].

In the course of compound screening, we discovered that unrelated classes of ATP-competitive dual EGFR/HER2 ligands, including thienopyrimidines [54] and thiazolylquinazolines [53] also bound to TRIB2 *in vitro*. In contrast to destabilization (a feature of covalent TRIB2 ligands) these compounds stabilize TRIB2 *in vitro*, similar to lapatinib and the pyrrolo[3,2-*d*] pyrimidine EGFR/HER compound TAK-285, which can bind to HER2 in an active-like conformation [76]. This might be equivalent to the thermostable TRIB2 generated by compound binding or after Cys 104 Tyr substitution. Moreover, evidence for a stabilized TRIB2:lapatinib complex was also established in cell extracts using CETSA (Supplementary Figure 10). Of relevance, the ATP-competitive ligands lapatinib and TAK-285 did not induce TRIB2 degradation in cells. However, previous studies demonstrated that lapatinib can deprive HER2 of an interaction with the Hsp90-Cdc37 system, leading to time-dependent HER2 degradation at micromolar concentrations [77]. Interestingly, like HER2, the cellular mechanism by which TRIB2 stability is regulated is proteasome-based (Figure 4), and we speculate that an afatinib-induced conformational change might induce TRIB2 ubiquitination, or negatively regulate interaction with an unknown stabilizing factor(s), similar to the effects of Hsp90 inhibitors towards the stability of Cdc37 kinome clients [78–80]. Consistently, TRIB2 is itself regulated by ubiquitination [18, 65, 66], and future work will evaluate TRIB2 binding to small molecules with dynamic changes in TRIB2 ubiqutination.

### The future: Targeting TRIB2 in cancer and beyond

The finding that TRIB2 stabilizing and destabilizing cell permeable ligands can be discovered by simple DSF profiling is an important advance for the evaluation of compounds that target this, and other, catalytically-deficient pseudokinases. The interaction of TRIB2 with both covalent and non-covalent clinical ligands confirms our original hypothesis, which stated that the TRIB2 pseudokinase represents a *bona fide* drug target [5]. We found that unique Cys residues in TRIB2 make it vulnerable to known families of small molecule kinase inhibitors that were originally developed as nanomolar covalent inhibitors of the tyrosine kinases EGFR and HER2. Such dual-targeting suggests shared features between signaling-relevant conformations in the ATP-site of both TRIB2 and EGFR/HER2, which generates reciprocal compound interactions. Conserved Cys residues in the atypical TRIB2 C-helical region most likely represent an additional form of selectivity filter, which permits targeting of TRIB2 by covalent classes of EGFR/HER2 inhibitor at micromolar concentrations in cells. Based on this mechanism, a lack of equivalent Cys residues in other pseudokinases, including the related TRIB1 and TRIB3, likely prevents them from interaction with covalent ligands such as afatinib.

Using a chemical genetic approach, we provide strong evidence that afatinib induces on-target effects through TRIB2 stability and AKT signaling in human cancer cells. Importantly, cellular signalling modulated by afatinib was prevented by mutation of two unique TRIB2 Cys residues, confirming a meaningful TRIB2 drug interaction in cells. Drug interaction was also validated by DSF, MS, MST and by employing in-cell TSA (CETSA) approaches with exogenous TRIB2. Finally, we established that afatinib (and neratinib) exhibit sub-micromolar toxicity in the human AML model cell line U937, where they are 10-20 fold more effective at cell killing compared to equipotent EGFR/HER2 or EGFR inhibitors [72], which do not degrade TRIB2 but still block ERK activation. U937 cells have previously been shown to be hypersensitive to TRIB2 siRNA knock-down [65], strengthening the case that the TRIB2 pseudokinase is rate-limiting for cell survival in this context. Experimental ‘on-target’ effects of covalent TRIB2-destablising agents were confirmed using chemical genetics and a Cys-mutant TRIB2 pseudokinase allele, similar to classical ‘drug-resistance’ approaches developed for compound target validation in the kinase inhibitor field [81–84].

## CONCLUSION

Our work demonstrates that covalent EGFR/HER2 inhibitors such as afatinib possess TRIB2 degrading-activity in human cells at micromolar concentrations, in a similar range to those reported for other destabilizing kinase inhibitors, including the covalent EGFR/HER2 drug neratinib [85]. Although we cannot rule out the simultaneous dual effects of afatinib on both TRIB2 and ERK/AKT phosphorylation contributing to cellular phenotypes, we provide clear evidence that TRIB2-binding is required for TRIB2 destabilization and AKT regulation in cells. Should an appropriate concentration of drug permit a direct TRIB2 interaction, an ‘off-target’ TRIB2-dependent phenotype might therefore be relevant to compound efficacy (or drug side-effects) in patient groups exposed to covalent EGFR/HER2 inhibitors. Many severe side effects of drugs are only detected after long-term clinical use, potentially leading to their withdrawal [86], and covalent drugs have the potential to accumulate to relatively high concentrations in cells. Interestingly, toxic side-effects such as diarrhea and vomiting induced by the EGFR/HER2 inhibitors lapatinib [87] and afatinib [88] are well-established in clinical patient cohorts. Based on the mechanistic studies reported here, it will be interesting to develop ELISA-based procedures to quantify effects of these drugs on TRIB2 protein stability in clinical samples obtained from drug-exposed patients, as part of broader proteomics approaches to establish all the intracellular targets of these compounds. Our study also provides impetus for generating improved (ideally TRIB2-specific) covalent ligands that induce TRIB2 degradation at even lower (sub-micromolar) concentrations, ideally by synthesising compounds in which the effects of eliminating or preserving EGFR/HER2 inhibition can be compared side-by-side. In the latter case, simultaneous elimination of TRIB2 and inhibition of the ERK-signaling pathway could be a polypharmacological asset, especially if TRIB2-dependent drug-resistance in tumor cells [38, 49] can be modulated by EGFR/HER2 inhibition. Taken together, our data establish a new paradigm for the pharmacological evaluation of agents that interfere with TRIB2-based signaling, and raise the intriguing possibility that dual EGFR/HER2 inhibitors might be repurposed as TRIB2-degrading agents in biological and clinical contexts. This information might also be exploited in the future for targeting a variety of TRIB2-overexpressing solid [89, 90] and haematological [20, 65] tumors.

## MATERIALS AND METHODS

### Chemicals, reagents, antibodies and TRIB2 small molecule screen

Tetracycline and doxycycline, MG132, AICAR and chloroquine were purchased from Sigma. Afatinib, neratinib, osimertinib, ibrutinib, erlotinib, lapatinib, TAK-285, BI2536, BI6727, gefitinib and bortezomib were purchased from LC laboratories or Selleck. Total AKT, pSer 473 AKT, pThr 308 AKT, total ERK1/2, dual pThr 202/pTyr 204 ERK1/2, cleaved Caspase 3 and α-tubulin antibodies were purchased from New England Biolabs and employed as previously described [50, 91]. 6His-HRP and α-FLAG antibodies were purchased from Sigma, GAPDH antibody was purchased from Proteintech and a polyclonal rabbit α-TRIB2 antibody was raised towards a unique N-terminal human TRIB2 sequence and affinity purified prior to evaluation with recombinant TRIB2 and a variety of human cellular extracts.

The PKIS chemical library (designated as SB, GSK or GW compounds) comprises 367 ATP-competitive kinase inhibitors, covering ∼30 chemotypes (∼70% with molecular mass <500 Da and clogP values <5) that were originally designated as ATP-competitive inhibitors of 24 distinct protein kinase targets, including multiple EGFR and HER2 tyrosine kinase classes [51]. Compounds were stored frozen as 10 mM stocks in DMSO. For initial screening, compounds were pre-incubated with TRIB2 for 10 minutes and then employed for DSF, which was initiated by the addition of fluorescent SYPRO Orange. For dose-dependent thermal-shift assays a compound range was prepared by serial dilution in DMSO, and added directly into the assay to the appropriate final concentration. All control experiments contained 2% (v/v) DMSO, which had essentially no effect on TRIB2 stability.

### Cloning, Site Directed Mutagenesis and recombinant protein production

pET30 6His-TRIB2 (and various deletion or amino acid substitution constructs, including an N-terminal FLAG-tagged TRIB2, termed His-FLAG-TRIB2), and pET30a 6His-PKAc, which encodes catalytically-active cAMP-dependent protein kinase domain, have been described previously [39, 48]. Full length TRIB2 was also cloned into pOPINJ to generate a His-GST-TRIB2 encoding construct for GST pull-down assays. Protein expression was induced in BL21(DE3) pLysS bacteria with 0.4 mM IPTG, and after overnight growth at 18°C, proteins were purified to near homogeneity using an initial affinity step (immobilised metal affinity chromatography (IMAC) or glutathione-sepharose chromatography) followed by size exclusion chromatography (16/600 Superdex 200) in appropriate buffers. Ser 473 Asp 6His-AKT1 (DU1850, amino acids 118-470), either active (PDK1-phosphorylated to a specific activity of 489 U/mg) or inactive, were purchased from the DSTT (University of Dundee), and stored at −80°C prior to analysis. Site directed mutagenesis was performed as previously described [92], using KOD Hot Start DNA polymerase and appropriate mutagenic primer pairs (sourced from IDT). All plasmids were Sanger-sequenced across the entire coding regions to confirm expected codon usage.

### Differential Scanning Fluorimetry (DSF)

Thermal-shift assays were performed using an Applied Biosystems StepOnePlus Real-Time PCR instrument using a standard DSF procedure previously developed and validated for the analysis of kinases [46, 48] and pseudokinases [47]. Proteins were diluted in 20 mM Tris/HCl (pH 7.4), 100 mM NaCl and 1 mM DTT to a concentration of 5 μM and then incubated with the indicated concentration of compound in a total reaction volume of 25 μL, with final concentration of 2% (v/v) DMSO. SYPRO Orange (Invitrogen) was used as a fluoresecence probe. The temperature was raised in regular 0.3 °C intervals from 25°C to 95°C. Compound binding experiments were assessed in duplicate and then reported relative to DMSO controls.

### Mass Spectrometry analysis of TRIB2 afatinib binding

To evaluate TRIB2 binding *in vitro*, afatinib was incubated for 15 min with purified 6His-TRIB2 at a 1:10 molar ratio, then denatured with 0.05 % (w/v) RapiGest SF (Waters, UK) and digested with chymotrypsin (1:20 protease:protein (w/w) ratio) for 16 h at 25 °C. RapiGest hydrolysis was induced by the addition of triflouroacetic acid (TFA) to 1 % (v/v), incubated at 37 °C for 1h. Insoluble product was removed by centrifugation (13, 000 × g, 20 min). Reversed-phase HPLC separation was performed using an UltiMate 3000 nano system (Dionex) coupled in-line with a Thermo Orbitrap Fusion Tribrid mass spectrometer (Thermo Scientific, Bremen, Germany). 500 fmol digested peptides were loaded onto the trapping column (PepMap100, C18, 300 μm × 5 mm), using partial loop injection, for 7 min at a flow rate of 9 μL/min with 2% (v/v) MeCN, 0.1% (v/v) TFA and then resolved on an analytical column (Easy-Spray C18 75 μm × 500 mm, 2 μm bead diameter column) using a gradient of 96.2% A (0.1% (v/v) formic acid (FA)): 3.8% B (80% (v/v) MeCN, 0.1% (v/v) FA) to 50% B over 35 min at a flow rate of 300 nL/min. MS1 spectra were acquired over *m/z* 400 – 1500 in the orbitrap (60 K resolution at 200 *m/z*). Data-dependent MS2 analysis was performed using a top speed approach (cycle time of 3 s), using higher-energy collisional dissociation (HCD) and electron-transfer and higher-energy collision dissociation (EThcD) for fragmentation, with product ions being detected in the ion trap (rapid mode). Data were processed using Thermo Proteome Discoverer (v. 1.4) and spectra were searched in MASCOT against the *E. coli* IPI database with the added sequence of full-length 6His-TRIB2 (1-343). Parameters were set as follows: MS1 tolerance of 10 ppm, MS/MS mass tolerance of 0.6 Da, oxidation of methionine and afatanib binding at cysteine as variable modifications. MS2 spectra were interrogated manually.

To evaluate the interaction between afatinib and intact TRIB2 protein, TRIB2 was incubated with afatinib as above, and then desalted using a C4 desalting trap (Waters MassPREP^TM^ Micro desalting column, 2.1 × 5 mm, 20 μ m particle size, 1000 Å pore size). TRIB2 was eluted with 50 % (v/v) MeCN, 0.1 % (v/v) formic acid. Intact TRIB2 mass analysis was performed using a Waters nano ACQUITY Ultra Performance liquid chromatography (UPLC) system coupled to a Waters SYNAPT G2. Samples were eluted from a C4 trap column at a flow rate of 10 μL/min using three repeated 0-100 % acetonitrile gradients. Data was collected between 400 and 3500 *m/z* and processed using MaxEnt1 (maximum entropy software, Waters Corporation).

### TRIB2 modeling and compound docking

A structural model of human TRIB2 kinase (UniProt ID: Q92519) containing the C-terminal tail (residues 42-343) was built, as described previously [12] using the crystal structure of TRIB1 (pdb: 5CEM) as the template. The N-terminal tail was not modelled since it was predicted to be disordered. The modeled TRIB2 structure was energy minimized using the Rosetta Relax protocol [93] and the model with the lowest energy score from 100 potential solutions was used for docking. The chemical structures of afatinib (CID 10184653) and neratinib (CID 9915743) were retrieved from PubChem.

Both compounds were independently docked using AutoDock Vina [62] with an exhaustiveness of 1000, rigid body, and a full protein search space. A binding pocket formed between P-loop, C-helix and the activation loop was the most frequently sampled site during blind docking. Compounds were docked to this putative docking site using the flexible side-chain covalent docking method implemented in Autodock [94]. In brief, the enamide group of each compound was covalently linked to the sulfhydryl group of Cys 96 and the rest of the compound was treated as a rigid side-chain to explore favorable binding poses, the vast majority of which converged on the same binding pocket. Each run generated 500 binding poses, which were visualized using PyMOL.

### MicroScale Thermophoresis

A Monolith NT.115 instrument (NanoTemper Technologies GmbH) was employed for MST analysis. His-TRIB2 was initially labelled with a NanoTemper labelling kit; the fluorescent red dye NT-647 was coupled via NHS chemistry to the N-terminal His-tag, placing the fluorophore away from the pseudokinase domain. For MST, the reaction was performed in 20 mM Bicine pH 9.0, 100 mM NaCl, 5% glycerol, 0.05% Tween-20 and 2 % (v/v) DMSO. Fluorescent TRIB2 (∼5 μM) was kept constant in the assay, while final afatinib concentration was titrated between 1 nM and 50 μM. Near-saturation binding was achieved, allowing for an affinity to be estimated for the reversible interaction of afatinib with the fluorescent TRIB2.

### Cell lines and reagents

Flp-In T-Rex parental HeLa cells (Invitrogen) were cultured in DMEM with 4 mM L-glutamine, 10 % (v/v) Foetal Bovine Serum (FBS), Penicillin and Streptomycin (Gibco) as described [81]. To engineer Tetracycline (Tet)-controlled expression of FLAG-tagged full length TRIB2 in human Flp-In T-REx cell lines, the host plasmid pcDNA5/FRT/TO, encoding full length TRIB2 sequences (or appropriate amino acid substitution(s)), with a single N-terminal FLAG tag (40 μg DNA per 10 cm dish of cells) were co-transfected with pOG44 Flp-Recombinase Expression Vector using lipofectamine. Cells that had successfully integrated the FLAG-tagged TRIB2 sequence were stably selected with 200 μg/mL Hygromycin B, according to the manufacturer’s instructions. Tet was added at a final concentration of 1 μg/ml to medium in order to induce FLAG-TRIB2 expression. For transient transfection, 50% confluent HeLa cells were transfected with 40 μg DNA per 10 cm dish for 48 h prior to lysis.

For serum starvation, stable HeLa cells were grown until ∼60% confluent in complete medium (+FBS), washed with PBS, and replaced with serum-free DMEM for 16 h. Cells were then incubated with DMEM containing 10% (v/v) FBS ± 1 μg/ml Tet for 16 h, followed by addition of appropriate inhibitor for 4h. Cells were then lysed, and whole cell extracts generated with modified RIPA buffer. Non-adherent AML-derived human U937 cells (which express high levels of endogenous TRIB2 protein) were supplied by Dr Karen Keeshan, University of Glasgow and were cultured as previously described [20].

### MTT cytotoxicity assay

U937 cells were seeded in a 96 well plate at a concentration of 0.2 × 10^6^ cells/mL, 18 hours prior to compound addition, which was performed in triplicate, with all experiments including a final concentration of 0.1% DMSO (v/v). To quantify U937 cell viability, metabolic activity was assessed 72 h after compound exposure using an MTT assay, as described previously [95]. Briefly, Thiazolyl blue tetrazolium bromide was dissolved in PBS and added to cells at a final concentration of 0.25 mg/mL and incubated at 37 °C for 3 hours. The reaction was stopped by the addition of 50 μL of acidified 10% SDS, followed by reading of absorbance at 570 nm. Viability was defined relative to DMSO-containing controls incubated for the same period of time.

### Immunoblotting and CETSA

HeLa and U937 whole cell lysates were generated by lysis in modified RIPA buffer (50 mM Tris-HCl pH 7.4, 1 % (v/v) NP-40, 1 % (v/v) SDS, 100 mM NaCl, 100 nM Okadaic acid) in the presence of protease and phosphatase inhibitors, and sonicated briefly to shear DNA prior to analysis. Samples were boiled for 5 min in sample buffer (50 mM Tris pH 6.8, 1% SDS, 10% glycerol, 0.01% Bromophenol Blue, 10 mM DTT). Subsequently, between 40 and 120 μg of total protein was resolved by SDS-PAGE followed by transfer onto nitrocellulose membrane (BioRad). After blocking in pH 7.4), primary and secondary antibodies were incubated in the same condition, and proteins were detected using HRP-conjugated antibodies and ECL reagent. Immunoblots were quantified using ImageJ software, as previously described [91]. For lambda phosphatase treatment, 40 μg FLAG-TRIB2 expressing stable cell extracts in lysis buffer without SDS and phosphatase inhibitors were incubated with 10 ng of purified λ-phosphatase for 30 min at 37°C prior to processing for western blotting.

For in-cell CETSA we employed a previously published procedure [96]. Briefly, stable HeLa cells were incubated with Tet for 16h to induce expression of FLAG-TRIB2, and at ∼90% confluence, were incubated with 0.1% (v/v) DMSO or 100 μM of the indicated compound for 1h. Intact cells were isolated by trypsinization (1 min) and resuspended in PBS, and then aliquoted into individual PCR tubes prior to heating at the indicated temperature in a PCR thermal cycler for 3 min. Cells were then placed on ice for 2 min and lysed by sonication, prior to centrifugation at 16,000 × g for 20 minutes at 4°C. The soluble lysate was analysed for the presence of FLAG-TRIB2 and α-tubulin by immunoblotting.

### Statistical analysis

All experimental procedures were repeated in at least 3 separate experiments (unless stated otherwise) and data are expressed as the mean ± standard deviation, where appropriate. When applied, statistical significance was defined as a p-value of ≤0.05, either using either a Students t-test or analysis of variance (ANOVA).

## ACKNOWLEDGEMENTS

The authors acknowledge Sam Evans for excellent technical support, especially media preparation.

## FUNDING

This work was funded by two BBSRC DTP studentships (to DMF and SF), a Tools and Resources Development Fund award (BB/N021703/1) to PAE, Royal Society Research Grants (to PAE and CEE), and North West Cancer Research grants (to PAE, CR1088 and CR1097). The SGC is a registered charity (number 1097737) that receives funds from AbbVie, Bayer Pharma AG, Boehringer Ingelheim, Canada Foundation for Innovation, Eshelman Institute for Innovation, Genome Canada, Innovative Medicines Initiative (EU/EFPIA) [ULTRA-DD grant no. 115766], Janssen, Merck KGaA Darmstadt Germany, MSD, Novartis Pharma AG, Ontario Ministry of Economic Development and Innovation, Pfizer, São Paulo Research Foundation-FAPESP, Takeda, and The Wellcome Trust [106169/ZZ14/Z].

## AUTHOR CONTRIBUTIONS

PAE obtained funding and designed experiments alongside DMF, DPB, FPB, SF and CEE. KK provided critical reagents and experimental advice. SS, WY and NK conducted TRIB2 and ligand modeling and docking. PAE generated the TRIB2 antibody, SF and CEE provided MS expertise; CW, DHD and WJZ provided the PKIS library, screening advice and additional medicinal chemistry. PAE wrote the paper, with contributions from all the authors, who approved the final version.

## COMPETING INTERESTS

There are no perceived conflicts of interest. The SGC receives direct funds from a variety of pharmaceutical companies (see above), although it remains entirely independent.

## DATA AND MATERIALS AVAILABILITY

Original TRIB2 PKIS screening data is available in Supplementary Table 1. Further compound information can be obtained by contacting DHD or WJZ.

**Supplementary Figure S1.**
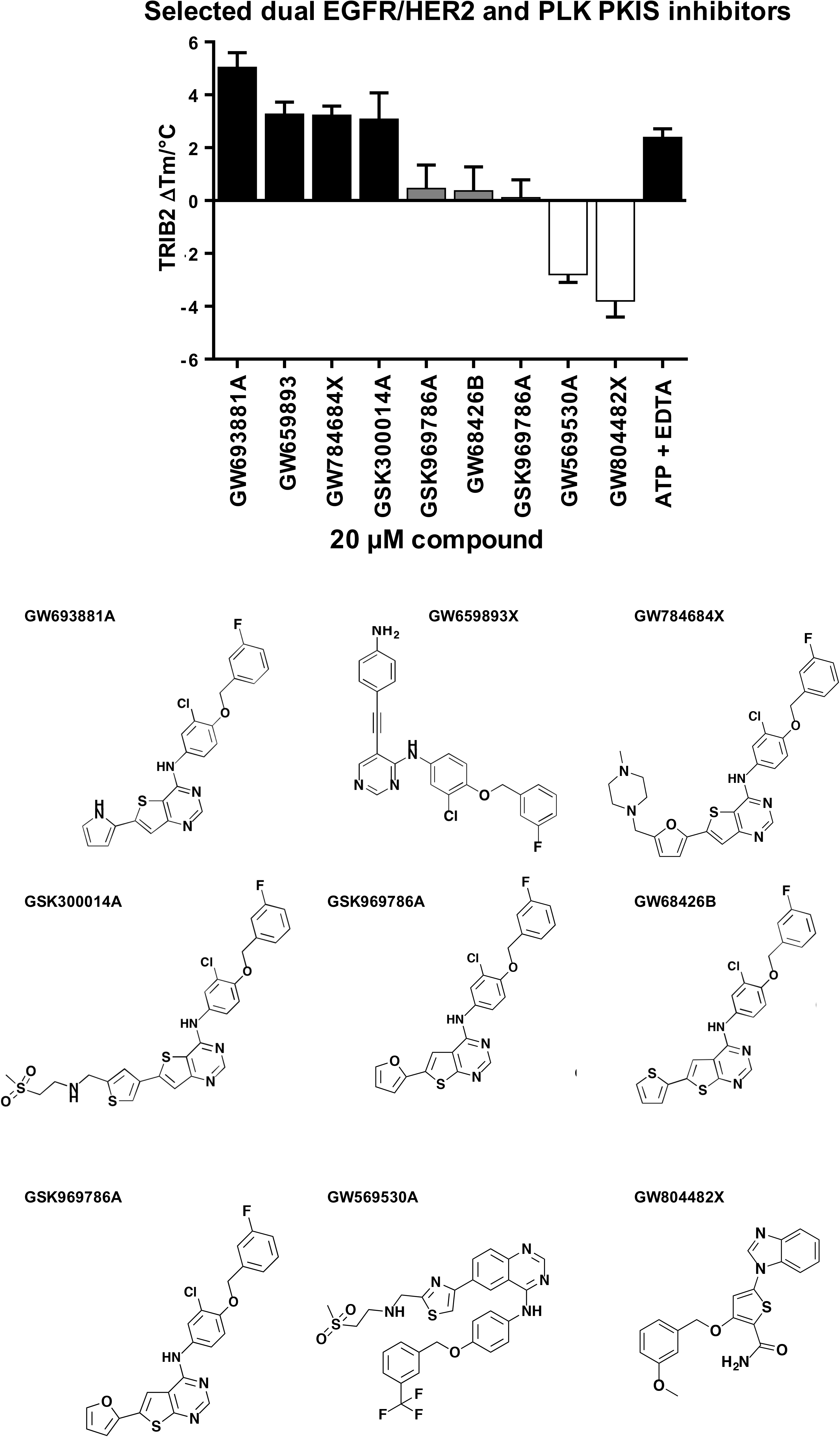
Discovery of multiple PKIS compounds as TRIB2 ligands. Dual EGFR/HER2 kinase inhibitors from three distinct chemical chemotypes present in the PKIS library were analysed by DSF using His-TRIB2 in a thermal shift assay. TRIB2 was screened with a 4-fold molar excess of each compound (20 μM). ΔT_m_ values were calculated for each condition, determined from fitting the Boltzmann equation. 10 mM ATP with 1mM EDTA (stabilization) and the TRIB2 thiophene ligand GW804482X (destabilization) were employed as controls. The chemical structures of class-specific PKIS compounds are also presented.

**Supplementary Figure S2.**
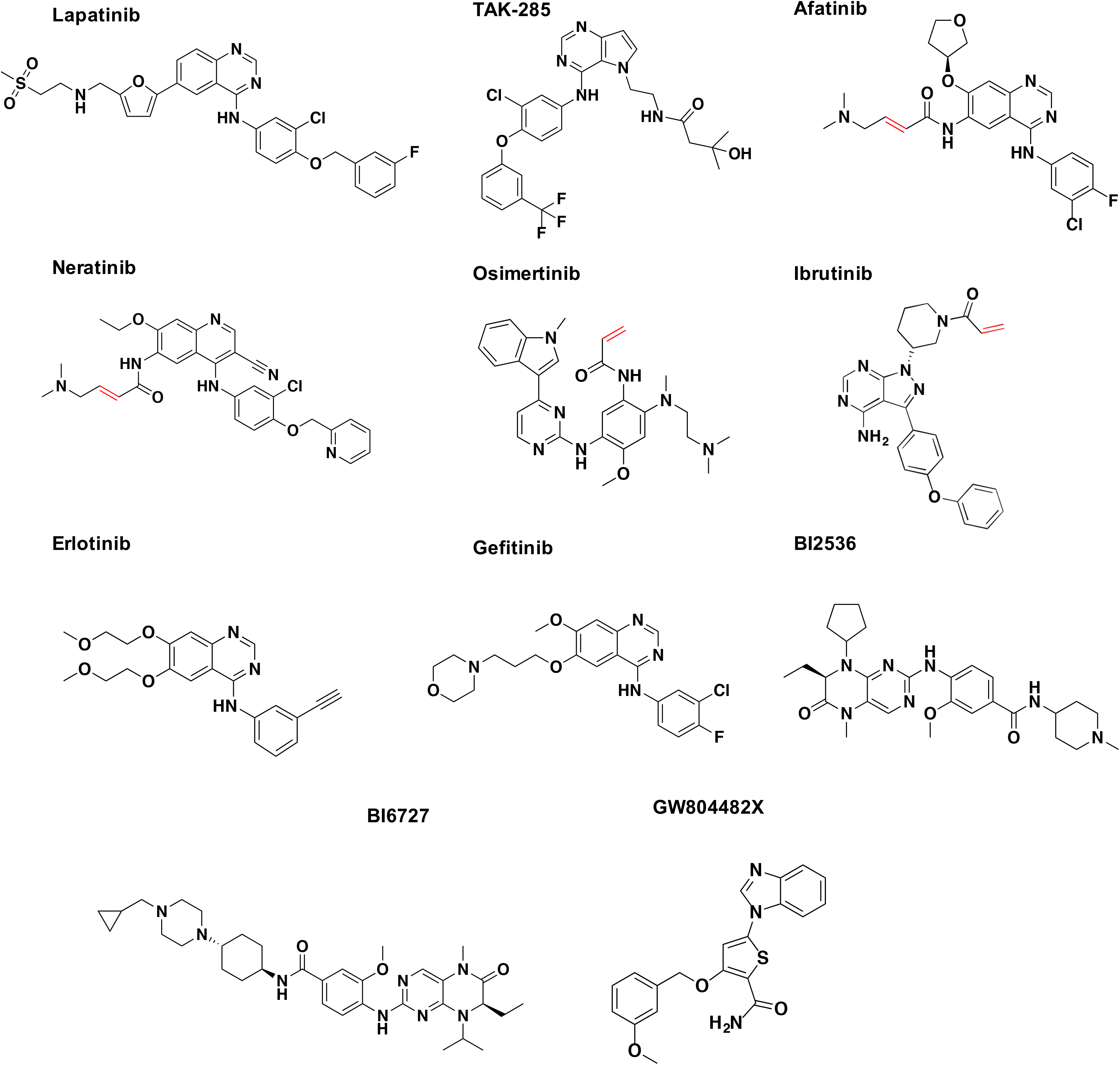
Chemical structure of pre-clinical and clinical compounds evaluated in this study. Eleven compounds are shown, including *in vitro* TRIB2 stabilizing agents such as lapatinib and TAK-285 and TRIB2 destabilizing compounds, such as afatinib, neratinib and osimertinib, all of which have previously been designated as dual EGFR/HER2 inhibitors. For covalent compounds, the electrophilic acrylamide group is shown in red. Erlotinib and gefitinib are potent (nM) clinical EGFR inhibitors, whereas BI2536 and BI6727 are potent (nM) PLK1-3 inhibitors. The thiophene TRIB2 destabilizer GW804482X (a PLK inhibitor) is included for comparison.

**Supplementary Figure S3.**
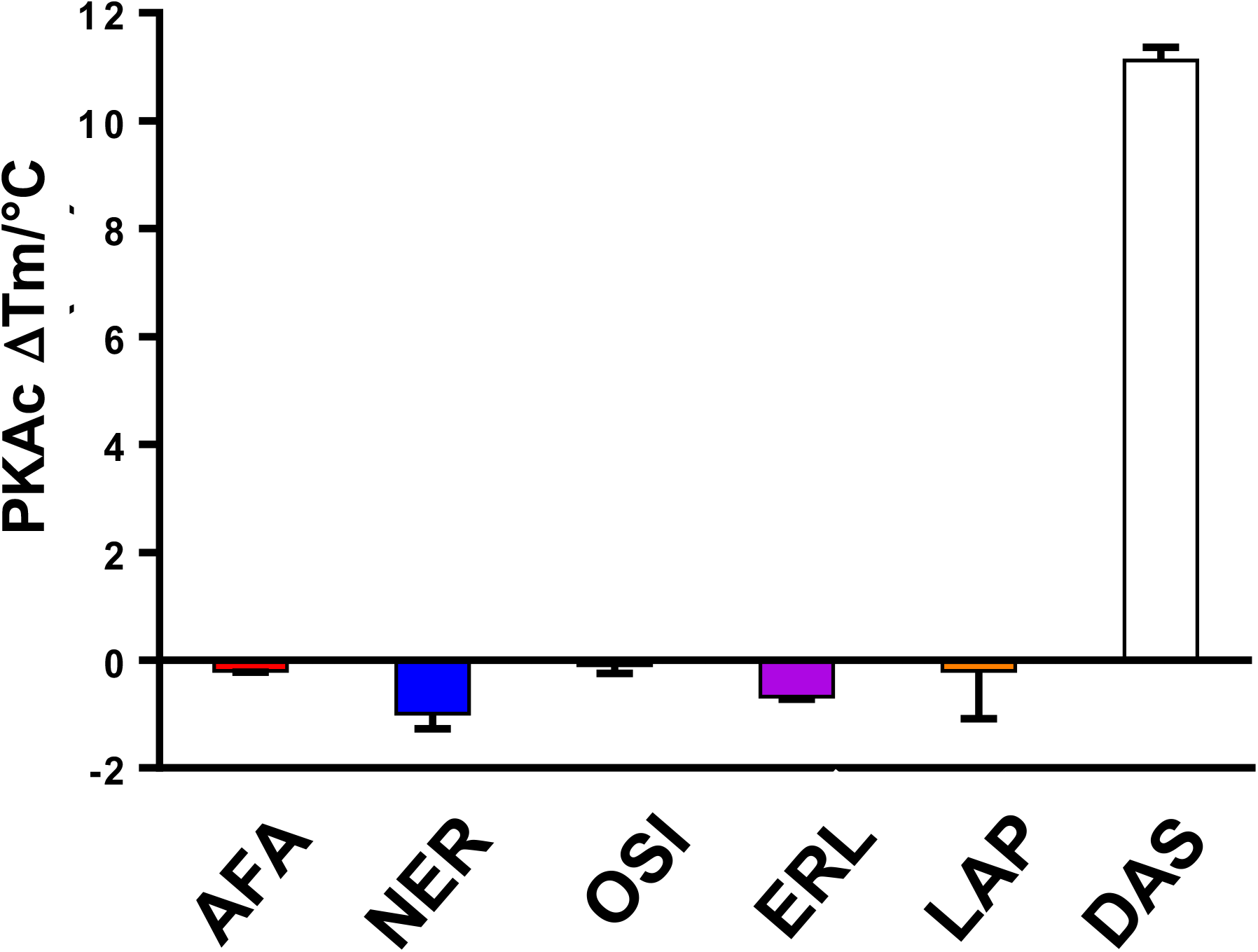
Validation of DSF assay using a PKAc counter-screen. Recombinant His-PKAc stability was analysed in the presence of a panel of dual EGFR/HER2 or EGFR inhibitors (final concentration 160 μM), confirming a lack of binding. The PKA stabilizing compound dasatinib (160 μM) was employed as a positive control. Data from two experiments (±SD) for triplicate assays are shown.

**Supplementary Figure S4.**
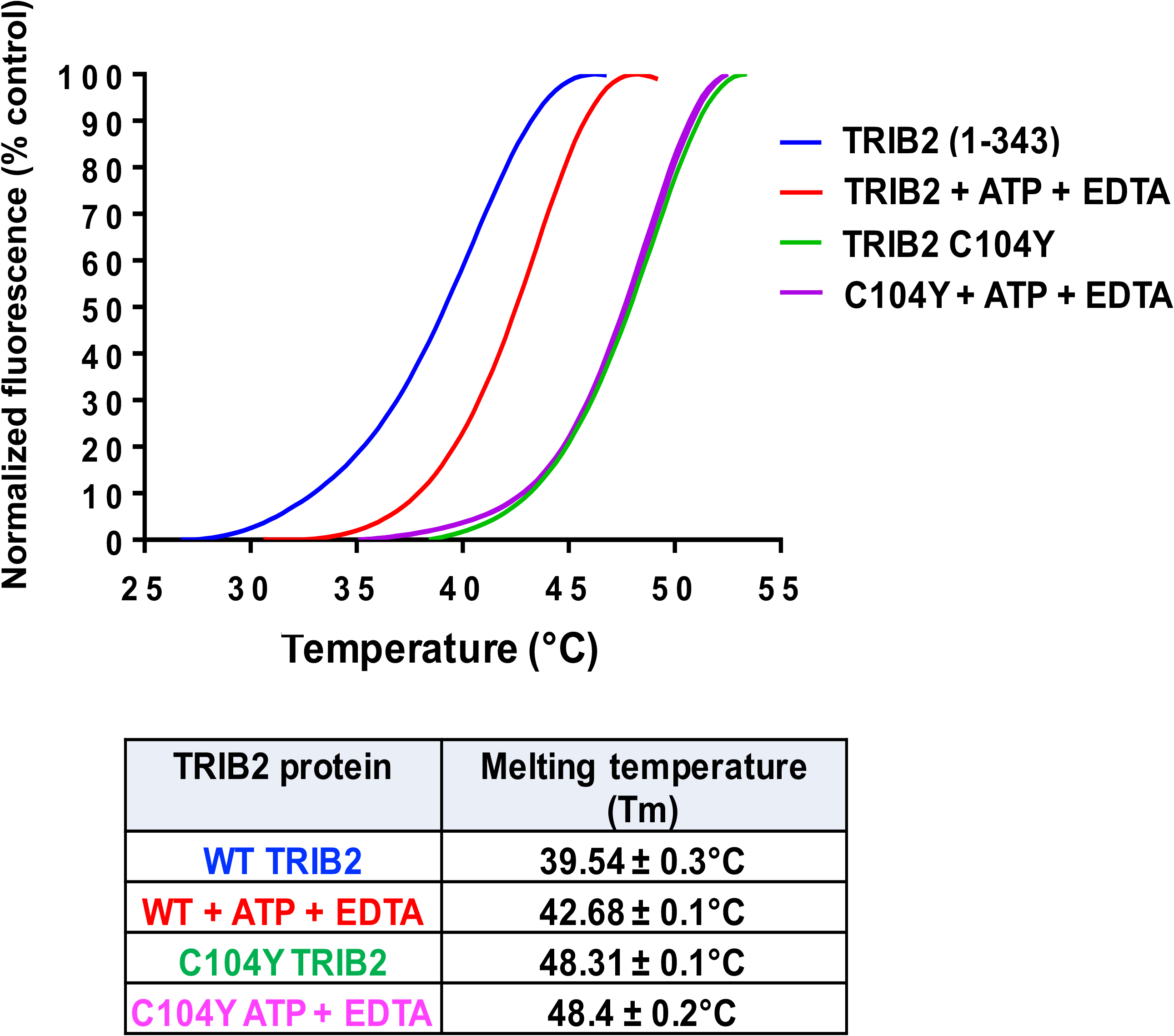
Loss of ATP-binding induced by C104Y mutation in TRIB2. Thermal denaturation profiles of full length WT-TRIB2 (1-343) or C104Y TRIB2 (1-343) proteins assessed in the presence or absence of 10 mM ATP, 1mM EDTA. A representative unfolding profile is shown for each. T_m_ values (±SD) were calculated from 3 separate experiments, each assayed in duplicate.

**Supplementary Figure S5.**
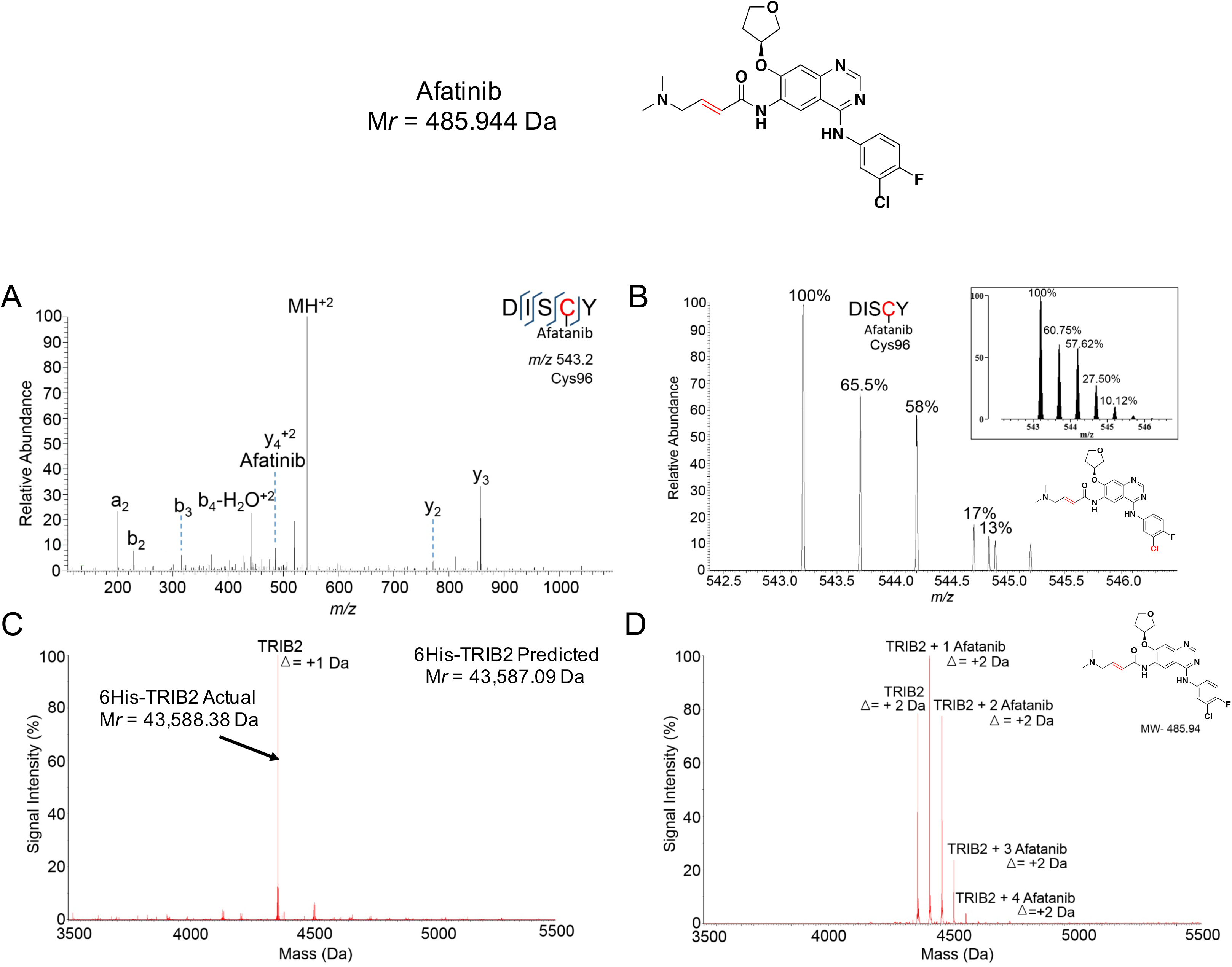
MS-based analysis of the covalent TRIB2:afatinib complex. MS2 spectra generated by HCD of chymotryptic peptides derived from 5 μM full length His-tagged TRIB2 (1-343) incubated with 20 μM afatinib for 15 minutes. (A) The doubly charged ion at *m/z* 543.2 confirms covalent binding of afatinb to Cys96 in the diagnostic Asp-Ile-Ser-Cys^96-^Tyr tetrapeptide derived from chymotrypsin. (B) The isotopic distribution for the 543.2 m/z ion precisely matches the theoretical isotopic distribution for an ion containing chlorine isotopes ^35^Cl and ^37^Cl, which are present in the spectrum due to the chlorofluro-benzyl moiety present in afatinib. (C) Intact mass spectrum for purified His-TRIB2, indicating a calculated molecular mass of 43,588.38 Da, compared to a predicted mass of 43,587.09. (D) Intact mass spectrum for purified TRIB2 preincubated with afatinib for 15 minutes at 20°C. The TRIB2 spectrum was dominated by peaks corresponding to intact TRIB2 covalently bound to either 0, 1 or 2 afatinib molecules, although up to 4 molecules of afatinib could be detected, with the number of covalent afatinib molecules increasing as a function of TRIB2 exposure time.

**Supplementary Figure S6.**
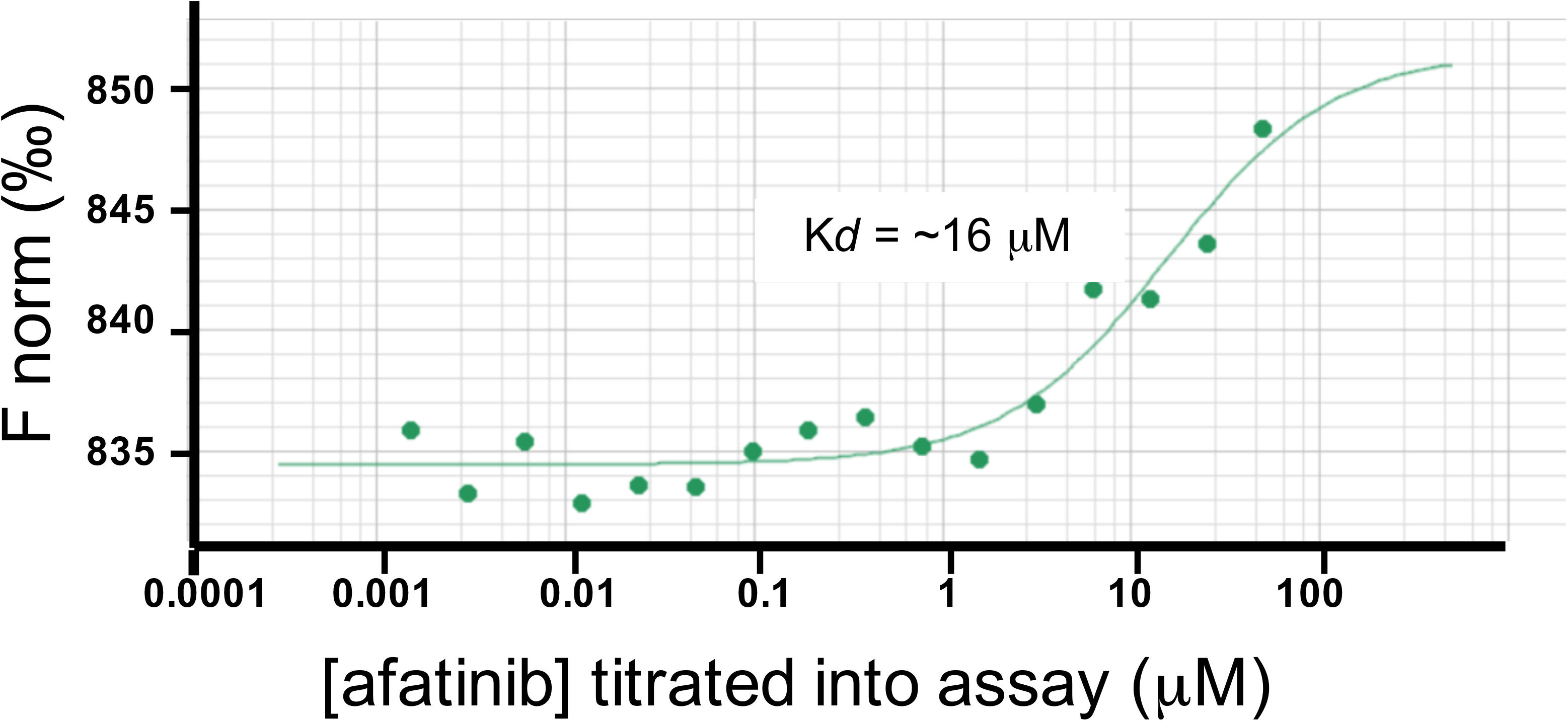
Microscale Thermophoresis (MST) assay. MST analysis of the His-TRIB2 and afatinib interaction in solution demonstrates a K*d* value of ∼16 μM for non-covalent reversible binding, a pre-requisite for formation of a covalent bond.

**Supplementary Figure S7.**
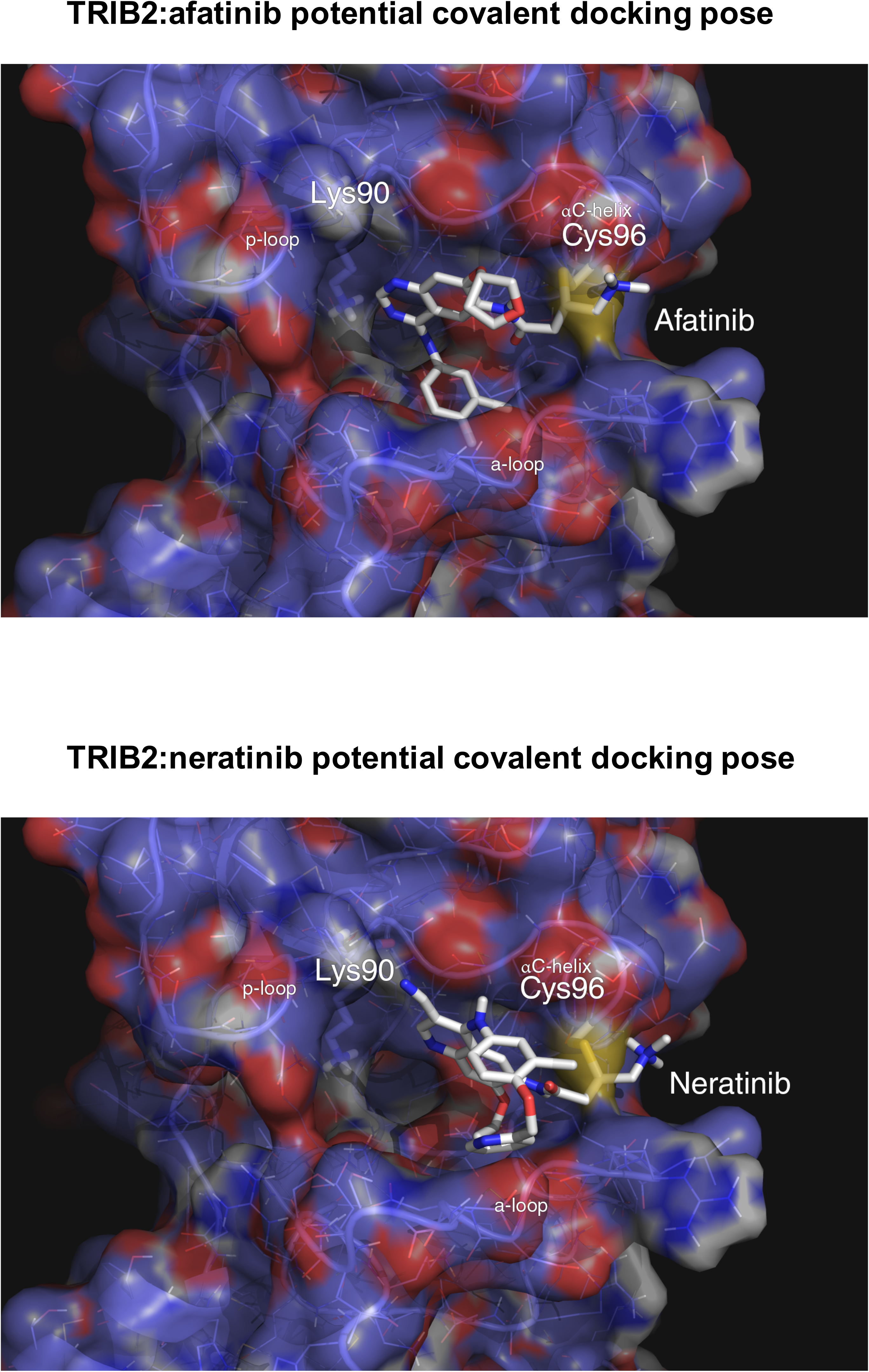
Molecular docking analysis. TRIB2 binding pose solutions for covalently bound afatinib or neratinib were visualized using PyMOL. The positions of Cys96 relative to the conserved β3 Lys (Lys 90) in TRIB2 are highlighted.

**Supplementary Figure S8.**
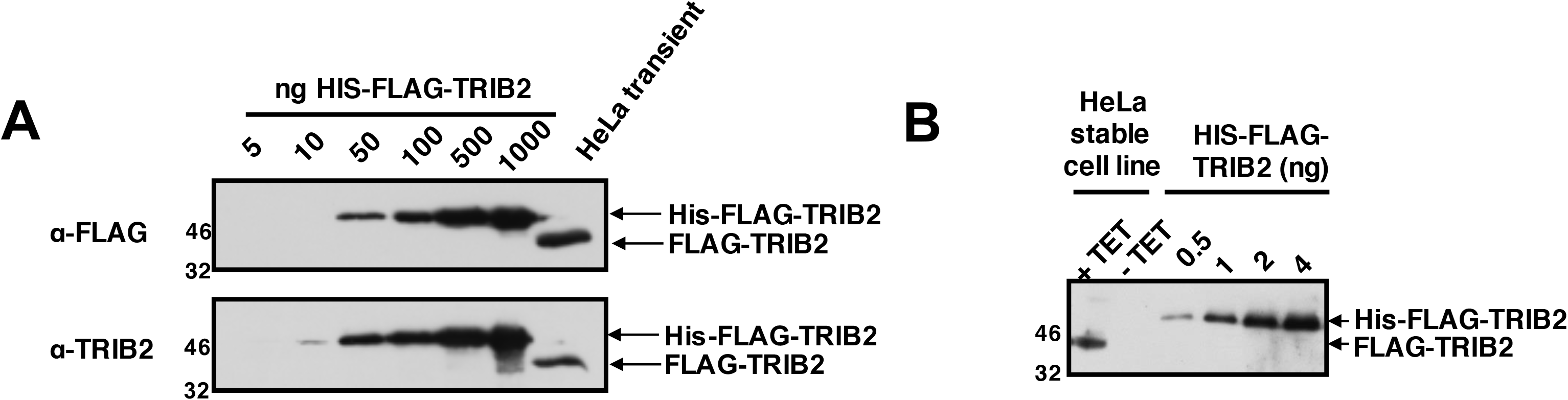
A new TRIB2 antibody for quantitative analysis of TRIB2 expression levels and stability. **(A)** The indicated amount of purified His-FLAG-TRIB2 and a HeLa cell extract containing transiently expressed FLAG-TRIB2 were used for immunoblotting with FLAG antibody (top), or side-by-side comparison with rabbit polyclonal TRIB2 antibody (bottom). **(B)** Analysis of FLAG-TRIB2 expression levels in stable HeLa cells in the presence or absence of Tetracycline (Tet). Known amounts of His-FLAG-TRIB2 were employed for quantification using FLAG antibody.

**Supplementary Figure S9.**
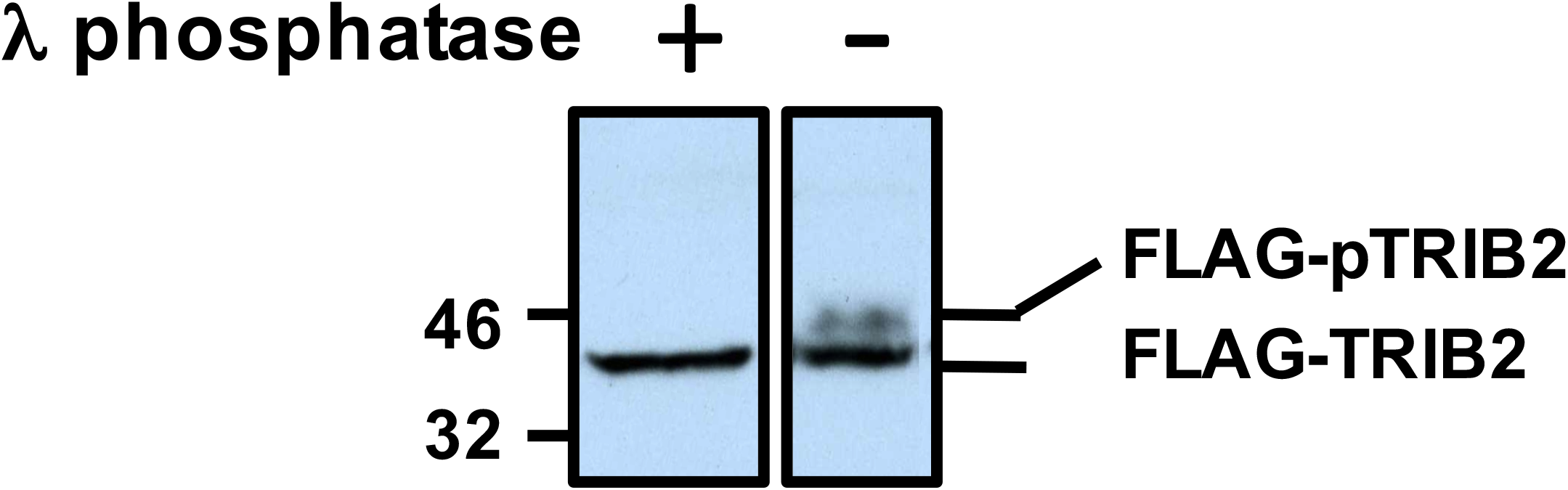
Analysis of TRIB2 dephosphorylation in cell extracts. FLAG-TRIB2 expression was induced with Tet for 16h, and 40 μg of appropriate whole cell lysates were treated with 10 ng lambda phosphatase for 1h revealing the presence of a slower-migrating, phosphorylated population of FLAG-TRIB2 in the cell extract.

**Supplementary Figure S10.**
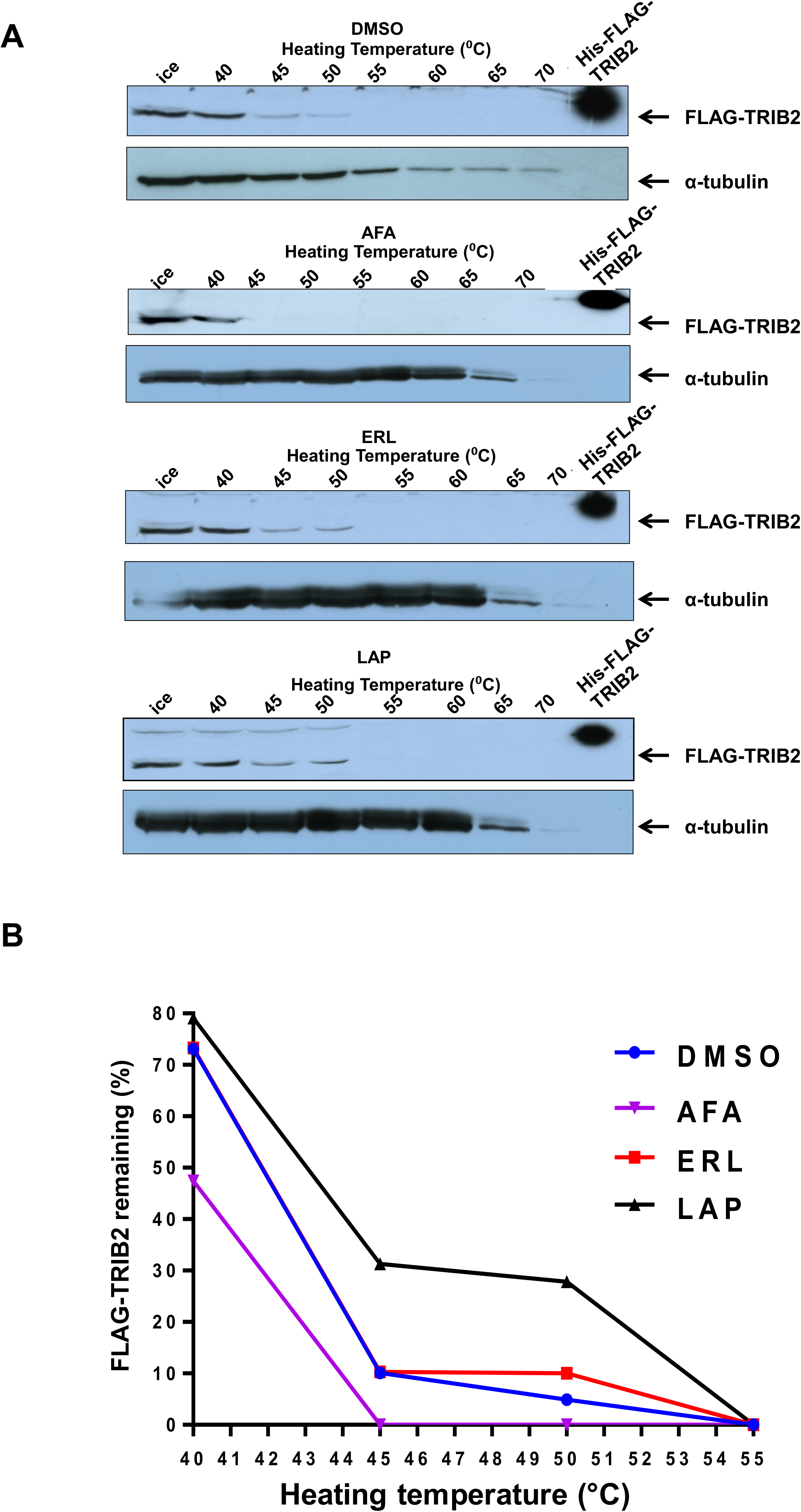
TRIB2-binding to afatinib induces destabilization relative to DMSO in a whole cell thermal shift assay (CETSA) **(A)** Cells were grown to 90% confluency and treated with either DMSO or 100 μM of the indicated compounds for 1 h. Cells were subsequently collected and re-suspended in PBS. The re-suspended cells were sub-aliquoted into PCR tubes and heated in a series of 5 °C increments for a total of 3 minutes at each temperature between 40 to 70 °C. The cells were subsequently lysed by sonication, centrifuged for 20 minutes at 18,000 g and analysed by western blot for either FLAG-TRIB2, alpha tubulin or 10 ng of His-FLAG-TRIB2. A representative experiment is shown **(B)** Densitometry analysis of CETSA experiment. TRIB2 levels (confirmed by gel migration relative to 10 ng recombinant His-FLAG-TRIB2) were normalised relative to unheated DMSO controls. Similar results were seen in an independent experiment.

**Supplementary Figure S11.**
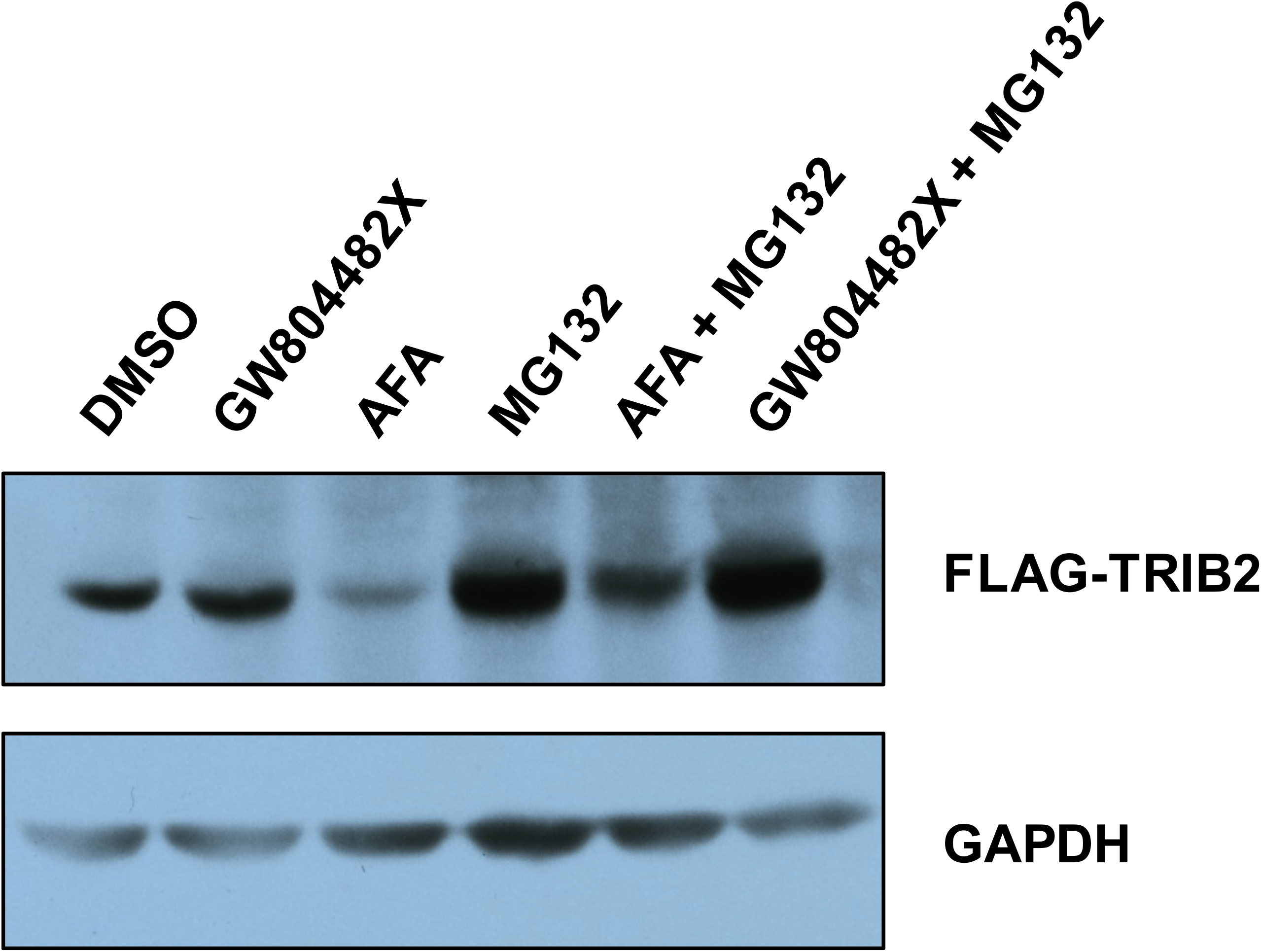
Lack of effect of the non-covalent TRIB2 destabilizing ligand GW804482X on TRIB2 stability in stable HeLa cells. FLAG-TRIB2-expression was induced with Tet for 24h, and cells were subsequently exposed for a further 4h to 10 μM GW804482X or 10 μM afatinib (AFA) in the presence or absence of 10 μM MG132. Cells were then lysed and processed for western blotting with FLAG or GAPDH antibodies.

**Supplementary Table 1.** PKIS compound screening data for full-length TRIB2. Data were obtained from duplicate DSF assays containing 5μM His-TRIB2 and 20 μM compound. n=2. Mean ΔT_m_ values reported ± SD.

| Plate position | Compound code | $\Delta T_m$ 1 | $\Delta T_m$ 2 | average | st.dev |
| --- | --- | --- | --- | --- | --- |
| JMP-A1 | SB-630812 | -0.3 | -0.27 | -0.285 | 0.015 |
| JMP-A2 | GW831091X | -0.09 | 0 | -0.045 | 0.045 |
| JMP-A3 | GW700494X | 2.83 | 2.06 | 2.445 | 0.385 |
| JMP-A4 | GW461104B | 1.2 | 1.51 | 1.355 | 0.155 |
| JMP-A5 | GW796920X | -0.12 | 0.07 | -0.025 | 0.095 |
| JMP-A6 | GW632046X | 0.6 | 0.43 | 0.515 | 0.085 |
| JMP-A7 | GW627834A | 1.27 | 1.71 | 1.49 | 0.22 |
| JMP-A8 | GSK561866B | -0.03 | 0.24 | 0.105 | 0.135 |
| JMP-A9 | GW352430A | 0.21 | 0.25 | 0.23 | 0.02 |
| JMP-A10 | SB-734117 | 0.63 | 0.64 | 0.635 | 0.005 |
| JMP-B1 | GW620972X | -0.93 | -0.19 | -0.56 | 0.37 |
| JMP-B2 | SB-686709-A | 0.1 | 0.13 | 0.115 | 0.015 |
| JMP-B3 | GW578748X | 0.29 | 0.13 | 0.21 | 0.08 |
| JMP-B4 | GW572399X | 1.81 | 2.06 | 1.935 | 0.125 |
| JMP-B5 | GSK2163632A | -2.96 | -0.38 | -1.67 | 1.29 |
| JMP-B6 | GSK953913A | -0.01 | 0.22 | 0.105 | 0.115 |
| JMP-B7 | GSK180736A | 0.27 | 0.32 | 0.295 | 0.025 |
| JMP-B8 | SB-264866 | 0.32 | 0.29 | 0.305 | 0.015 |
| JMP-B9 | GW809897X | 1.63 | 1 | 1.315 | 0.315 |
| JMP-B10 | SB-361058 | 0.44 | 0.47 | 0.455 | 0.015 |
| JMP-C1 | GW829055X | 1.43 | 1.68 | 1.555 | 0.125 |
| JMP-C2 | GW434756X | -0.12 | 0.11 | -0.005 | 0.115 |
| JMP-C3 | GSK237700A | -0.36 | -0.55 | -0.455 | 0.095 |
| JMP-C4 | GW577921A | 0.77 | 0.03 | 0.4 | 0.37 |
| JMP-C5 | GW621970A | 0.3 | 0.36 | 0.33 | 0.03 |
| JMP-C6 | SB-741905 | 0.06 | 0.22 | 0.14 | 0.08 |
| JMP-C7 | SB-736290 | 0.11 | 0.03 | 0.07 | 0.04 |
| JMP-C8 | SB-742865 | 1.01 | 0.83 | 0.92 | 0.09 |
| JMP-C9 | GW874091X | 0.27 | 0.26 | 0.265 | 0.005 |
| JMP-C10 | SB-738561 | -1.22 | 0.34 | -0.44 | 0.78 |
| JMP-D1 | GW829115X | 0.05 | 0.28 | 0.165 | 0.115 |
| JMP-D2 | GW856804X | 0.96 | 1.15 | 1.055 | 0.095 |
| JMP-D3 | GW833373X | 1.05 | 0.91 | 0.98 | 0.07 |
| JMP-D4 | GW796921X | -0.02 | 0.28 | 0.13 | 0.15 |
| JMP-D5 | GSK586581B | 0 | 0.21 | 0.105 | 0.105 |
| JMP-D6 | SB-390527 | 0.24 | 0.42 | 0.33 | 0.09 |
| JMP-D7 | GSK571989A | -0.33 | 0.05 | -0.14 | 0.19 |
| JMP-D8 | GW784684X | 3.58 | 2.88 | 3.23 | 0.35 |
| JMP-D9 | GSK204925A | 0.22 | 0.72 | 0.47 | 0.25 |
| JMP-D10 | SB-236687 | 0.45 | -0.93 | -0.24 | 0.69 |
| JMP-E1 | GW829906X | 0.52 | 1.29 | 0.905 | 0.385 |
| JMP-E2 | GW813360X | 0.04 | 0.15 | 0.095 | 0.055 |
| JMP-E3 | GW680908A | 1.37 | 1.33 | 1.35 | 0.02 |
| JMPE4 | SKF-62604 | -0.11 | -0.07 | -0.09 | 0.02 |
| JMP-E5 | GSK1023156A | 0.77 | 0.24 | 0.505 | 0.265 |
| JMP-E6 | SB-751399 | 0.16 | 0.6 | 0.38 | 0.22 |
| JMP-E7 | GW574783B | 1.59 | 2.69 | 2.14 | 0.55 |
| JMP-E8 | GW641155B | 0.52 | 0.52 | 0.52 | 0 |
| JMP-E9 | GW301789X | 0.89 | 0.95 | 0.92 | 0.03 |
| JMP-E10 | GW607049B | 3.48 | 2.93 | 3.205 | 0.275 |
| JMP-F1 | GSK620503A | 0.1 | -0.36 | -0.13 | 0.23 |
| JMP-F2 | GW569530A | -2.56 | -1.87 | -2.215 | 0.345 |
| JMP-F3 | SB-264865 | -2.97 | 0.2 | -1.385 | 1.585 |
| JMP-F4 | GSK319347A | 0.39 | 0.6 | 0.495 | 0.105 |
| JMP-F5 | SB-739452 | 0.47 | 0.4 | 0.435 | 0.035 |
| JMP-F6 | GSK711701A | 0.14 | 0.65 | 0.395 | 0.255 |
| JMP-F7 | GW768505A | 1.31 | 1.14 | 1.225 | 0.085 |
| JMP-F8 | SB-814597 | 0.35 | 0.46 | 0.405 | 0.055 |
| JMP-F9 | GW837331X | 3.16 | 1.88 | 2.52 | 0.64 |
| JMP-F10 | SB-678557-A | 0.57 | 0.5 | 0.535 | 0.035 |
| JMP-G1 | GW775608X | 0.12 | 0.32 | 0.22 | 0.1 |
| JMP-G2 | GI261520A | 1.58 | 0.81 | 1.195 | 0.385 |
| JMP-G3 | SB-245392 | -0.19 | 0.09 | -0.05 | 0.14 |
| JMP-G4 | GSK317315A | 0.43 | 0.23 | 0.33 | 0.1 |
| JMP-G5 | GW817396X | 0.08 | 0.38 | 0.23 | 0.15 |
| JMP-G6 | GW785804X | 0.1 | 0.43 | 0.265 | 0.165 |
| JMP-G7 | SB-732881-H | 0.03 | 0.41 | 0.22 | 0.19 |
| JMP-G8 | GSK1173862A | 1.27 | 1.6 | 1.435 | 0.165 |
| JMP-G9 | GW301888X | 0.53 | 0.64 | 0.585 | 0.055 |
| JMP-G10 | GW843682X | 0.66 | 0.42 | 0.54 | 0.12 |
| JMP-H1 | GW405841X | 1.25 | 1.24 | 1.245 | 0.005 |
| JMP-H2 | SB-409513 | 0.07 | 0.19 | 0.13 | 0.06 |
| JMP-H3 | GSK619487A | 0.03 | 0.07 | 0.05 | 0.02 |
| JMP-H4 | GW410563A | 0.07 | 0.27 | 0.17 | 0.1 |
| JMP-H5 | GW784752X | 0.27 | 0.4 | 0.335 | 0.065 |
| JMP-H6 | SKF-86002 | 0.25 | 0.32 | 0.285 | 0.035 |
| JMP-H7 | SB-376719 | 0.34 | 0.33 | 0.335 | 0.005 |
| JMP-H8 | SB-223133 | 0.24 | 0.29 | 0.265 | 0.025 |
| JMP-H9 | GW708893X | 0.54 | 0.04 | 0.29 | 0.25 |
| JMP-H10 | GW759710A | 0.79 | 0.7 | 0.745 | 0.045 |
| JOJ-A1 | GW416469X | -1.26 | -0.97 | -1.115 | 0.145 |
| JOJ-A2 | SKF-86055 | -0.71 | 0.09 | -0.31 | 0.4 |
| JOJ-A3 | SB-682330-A | -1.14 | -1.04 | -1.09 | 0.05 |
| JOJ-A4 | SB-242719 | -0.55 | -0.85 | -0.7 | 0.15 |
| JOJ-A5 | SB-333612 | 0.18 | -0.13 | 0.025 | 0.155 |
| JOJ-A6 | GSK1030061A | -0.41 | -0.95 | -0.68 | 0.27 |
| JOJ-A7 | GSK1030059A | -0.6 | -0.58 | -0.59 | 0.01 |
| JOJ-A8 | GW799251X | 1.05 | -0.19 | 0.43 | 0.62 |
| JOJ-A9 | GW832467X | 0.2 | 0.51 | 0.355 | 0.155 |
| JOJ-A10 | GW296115X | 2.67 | 3.85 | 3.26 | 0.59 |
| JOJ-B1 | GW830263A | -0.27 | -0.11 | -0.19 | 0.08 |
| JOJ-B2 | GSK1751853A | -0.32 | -0.43 | -0.375 | 0.055 |
| JOJ-B3 | GW643971X | 0.84 | 0.3 | 0.57 | 0.27 |
| JOJ-B4 | SB-614067-R | 0.48 | -0.55 | -0.035 | 0.515 |
| JOJ-B5 | GW694590A | 3.6 | 3.3 | 3.45 | 0.15 |
| JOJ-B6 | SB-278538 | -1.06 | -1.31 | -1.185 | 0.125 |
| JOJ-B7 | SB-390523 | -0.53 | -0.06 | -0.295 | 0.235 |
| JOJ-B8 | GSK200398A | -0.55 | -0.29 | -0.42 | 0.13 |
| JOJ-B9 | GW829877X | -0.19 | 0.17 | -0.01 | 0.18 |
| JOJ-B10 | GW435821X | 0.63 | 0.58 | 0.605 | 0.025 |
| JOJ-C1 | GSK938890A | -1.42 | -3.46 | -2.44 | 1.02 |
| JOJ-C2 | GW445015X | 0.1 | -0.52 | -0.21 | 0.31 |
| JOJ-C3 | GW853609X | 3.48 | 2.86 | 3.17 | 0.31 |
| JOJ-C4 | SB-400868-A | -0.44 | -0.56 | -0.5 | 0.06 |
| JOJ-C5 | SB-739245-AC | -0.25 | -0.88 | -0.565 | 0.315 |
| JOJ-C6 | SB-226879 | -0.65 | -0.72 | -0.685 | 0.035 |
| JOJ-C7 | GSK1030062A | -0.62 | -0.7 | -0.66 | 0.04 |
| JOJ-C8 | GW300657X | 0.35 | -0.26 | 0.045 | 0.305 |
| JOJ-C9 | SB-633825 | -1.34 | -2.17 | -1.755 | 0.415 |
| JOJ-C10 | GSK1030058A | 0.38 | 0.44 | 0.41 | 0.03 |
| JOJ-D1 | GW810372X | -0.31 | 0.57 | 0.13 | 0.44 |
| JOJ-D2 | GW627512B | -0.33 | 0 | -0.165 | 0.165 |
| JOJ-D3 | SB-360741 | -0.12 | -0.5 | -0.31 | 0.19 |
| JOJ-D4 | GW580509X | -0.16 | -0.25 | -0.205 | 0.045 |
| JOJ-D5 | GW631581B | 1.1 | -0.12 | 0.49 | 0.61 |
| JOJ-D6 | GSK1000163A | -3.2 | 0.18 | -1.51 | 1.69 |
| JOJ-D7 | GW703087X | 3.01 | 3.59 | 3.3 | 0.29 |
| JOJ-D8 | GW869810X | 1.31 | 1.88 | 1.595 | 0.285 |
| JOJ-D9 | GW780159X | -0.66 | -0.55 | -0.605 | 0.055 |
| JOJ-D10 | SB-239272 | 0.31 | -0.27 | 0.02 | 0.29 |
| JOJ-E1 | GW612286X | -2.41 | -0.83 | -1.62 | 0.79 |
| JOJ-E2 | GW284408X | -1.45 | -1.17 | -1.31 | 0.14 |
| JOJ-E3 | GW450241X | -1.05 | -0.9 | -0.975 | 0.075 |
| JOJ-E4 | GSK300014A | 2.08 | 4.08 | 3.08 | 1 |
| JOJ-E5 | GW770220A | -0.97 | -0.67 | -0.82 | 0.15 |
| JOJ-E6 | GW785974X | -0.56 | -0.72 | -0.64 | 0.08 |
| JOJ-E7 | GW828525X | 1.55 | 2.82 | 2.185 | 0.635 |
| JOJ-E8 | GW406108X | -0.66 | 1.04 | 0.19 | 0.85 |
| JOJ-E9 | GW804482X | -3.41 | -3.63 | -3.52 | 0.395 |
| JOJ-E10 | GSK299115A | 0.02 | 0.93 | 0.475 | 0.455 |
| JOJ-F1 | GW290597X | -0.76 | -0.81 | -0.785 | 0.025 |
| JOJ-F2 | GW644007X | -0.84 | 0.26 | -0.29 | 0.55 |
| JOJ-F3 | GW831090X | -0.17 | -0.05 | -0.11 | 0.06 |
| JOJ-F4 | SB-409514 | -0.72 | -0.56 | -0.64 | 0.08 |
| JOJ-F5 | GW794726X | -0.13 | 0.17 | 0.02 | 0.15 |
| JOJ-F6 | GW827099X | 0.98 | 1.08 | 1.03 | 0.05 |
| JOJ-F7 | GW827106X | -0.05 | 0.11 | 0.03 | 0.08 |
| JOJ-F8 | GW846105X | 0.7 | 1.77 | 1.235 | 0.535 |
| JOJ-F9 | SB-738482 | 0.94 | 1.32 | 1.13 | 0.19 |
| JOJ-F10 | GW827396X | 2.68 | 3.19 | 2.935 | 0.255 |
| JOJ-G1 | SB-253228 | -2.45 | -0.28 | -1.365 | 1.085 |
| JOJ-G2 | GW300653X | 0.52 | 0.45 | 0.485 | 0.035 |
| JOJ-G3 | GW695874X | -1.08 | 0.01 | -0.535 | 0.545 |
| JOJ-G4 | GW575533A | 1.31 | -0.2 | 0.555 | 0.755 |
| JOJ-G5 | SB-657836-AAA | -2.54 | -0.61 | -1.575 | 0.965 |
| JOJ-G6 | GW807930X | 1.98 | 1.81 | 1.895 | 0.085 |
| JOJ-G7 | GSK943949A | -1.1 | -1.77 | -1.435 | 0.335 |
| JOJ-G8 | GW659386A | 2.33 | 3.11 | 2.72 | 0.39 |
| JOJ-G9 | GSK1511931A | -0.17 | -0.1 | -0.135 | 0.035 |
| JOJ-G10 | GSK969786A | 1.54 | 1.76 | 1.65 | 0.11 |
| JOJ-H1 | GW568326X | -2.48 | -0.07 | -1.275 | 1.205 |
| JOJ-H2 | GW861893X | 1.42 | 1.89 | 1.655 | 0.235 |
| JOJ-H3 | GW275944X | 2.5 | 1.23 | 1.865 | 0.635 |
| JOJ-H4 | GW278681X | 0.19 | -0.58 | -0.195 | 0.385 |
| JOJ-H5 | SB-711237 | -0.15 | 1.12 | 0.485 | 0.635 |
| JOJ-H6 | GW305178X | 0.1 | 1.6 | 0.85 | 0.75 |
| JOJ-H7 | GW445017X | -0.06 | -0.61 | -0.335 | 0.275 |
| JOJ-H8 | GW440139B | -0.65 | -0.52 | -0.585 | 0.065 |
| JOJ-H9 | GW651576X | -0.19 | -0.55 | -0.37 | 0.18 |
| JOJ-H10 | GW566221B | 1.96 | 1.44 | 1.7 | 0.26 |
| JOK-A1 | GSK182497A | 1.21 | 1.4 | 1.305 | 0.095 |
| JOK-A2 | GW778894X | -0.83 | -0.52 | -0.675 | 0.155 |
| JOK-A3 | SB-220455 | -1.1 | -1.4 | -1.25 | 0.15 |
| JOK-A4 | GW429374A | -0.16 | 0.05 | -0.055 | 0.105 |
| JOK-A5 | GW784307A | -0.54 | -0.52 | -0.53 | 0.01 |
| JOK-B1 | GW781673X | -1.53 | -0.99 | -1.26 | 0.27 |
| JOK-B2 | SB-592602 | -1.13 | -0.7 | -0.915 | 0.215 |
| JOK-B3 | SB-278539 | -1.25 | -1.24 | -1.245 | 0.005 |
| JOK-B4 | GSK2213727A | -0.73 | -0.77 | -0.75 | 0.02 |
| JOK-B5 | GW576484X | 1.97 | 2.06 | 2.015 | 0.045 |
| JOK-C1 | GSK238583A | -1.63 | -1.19 | -1.41 | 0.22 |
| JOK-C2 | GSK1392956A | -0.89 | -1.23 | -1.06 | 0.17 |
| JOK-C3 | SB-437013 | -1.59 | -1.55 | -1.57 | 0.02 |
| JOK-C4 | SB-242717 | -0.72 | -0.91 | -0.815 | 0.095 |
| JOK-D1 | GW827105X | 1.94 | 2.08 | 2.01 | 0.07 |
| JOK-D2 | GW830365A | -1.76 | -0.93 | -1.345 | 0.415 |
| JOK-D3 | GSK317354A | -0.48 | -0.86 | -0.67 | 0.19 |
| JOK-D4 | SB-742864 | -0.93 | -0.88 | -0.905 | 0.025 |
| JOK-E1 | GW441756X | -1.97 | -1.17 | -1.57 | 0.4 |
| JOK-E2 | GR105659X | -1.04 | -1.13 | -1.085 | 0.045 |
| JOK-E3 | SB-735465 | -1.06 | -0.99 | -1.025 | 0.035 |
| JOK-E4 | GW806742X | 2.59 | 2.69 | 2.64 | 0.05 |
| JOK-F1 | GW589933X | -0.75 | -1.04 | -0.895 | 0.145 |
| JOK-F2 | SB-750140 | -0.8 | -0.92 | -0.86 | 0.06 |
| JOK-F3 | GW549390X | -0.08 | 0.29 | 0.105 | 0.185 |
| JOK-F4 | GW458344A | -1.7 | -1.26 | -1.48 | 0.22 |
| JOK-G1 | GW810576X | -1.23 | -0.69 | -0.96 | 0.27 |
| JOK-G2 | GW772405X | -0.13 | 0.62 | 0.245 | 0.375 |
| JOK-G3 | GW282974A | -0.64 | -0.2 | -0.42 | 0.22 |
| JOK-G4 | SB-431533 | -0.37 | -0.8 | -0.585 | 0.215 |
| JOK-H1 | GW853606X | 0.82 | 0.72 | 0.77 | 0.05 |
| JOK-H2 | SB-732941 | -0.67 | -0.83 | -0.75 | 0.08 |
| JOK-H3 | GW581744X | -1.13 | -0.58 | -0.855 | 0.275 |
| JOK-H4 | SB-221466 | -0.53 | -0.41 | -0.47 | 0.06 |
| JOI-A1 | GW673715X | 1.75 | 2.32 | 2.035 | 0.285 |
| JOI-A2 | GW811761X | 1.02 | -0.2 | 0.41 | 0.61 |
| JOI-A3 | GW680975X | 0.89 | -0.02 | 0.435 | 0.455 |
| JOI-A4 | GSK605714A | 0.88 | 0.3 | 0.59 | 0.29 |
| JOI-A5 | GW782612X | -0.01 | 0.37 | 0.18 | 0.19 |
| JOI-B1 | GSK192082A | 2.74 | 1.72 | 2.23 | 0.51 |
| JOI-B2 | SB-707548-A | 4.01 | 3.55 | 3.78 | 0.23 |
| JOI-B3 | SB-285234-W | 0.81 | -0.27 | 0.27 | 0.54 |
| JOI-B4 | GR269666A | -0.27 | 0.14 | -0.065 | 0.205 |
| JOI-B5 | GW559768X | 0.7 | 0.38 | 0.54 | 0.16 |
| JOI-C1 | SB-675259-M | -0.32 | -0.21 | -0.265 | 0.055 |
| JOI-C2 | GSK718429A | 3.26 | 2.35 | 2.805 | 0.455 |
| JOI-C3 | GW572738X | 0.9 | -0.11 | 0.395 | 0.505 |
| JOI-C4 | GW621431X | 0.64 | -0.09 | 0.275 | 0.365 |
| JOI-C5 | GW416981X | 0.97 | 0.39 | 0.68 | 0.29 |
| JOI-D1 | GSK237701A | 0.54 | -1.87 | -0.665 | 1.205 |
| JOI-D2 | GW567808A | 2.72 | 2.37 | 2.545 | 0.175 |
| JOI-D3 | GSK605714A | 0.76 | -0.01 | 0.375 | 0.385 |
| JOI-D4 | GW561436X | 0.78 | -0.04 | 0.37 | 0.41 |
| JOI-D5 | GW607117X | 0.61 | 0.7 | 0.655 | 0.045 |
| JOI-E1 | GSK978744A | 0.64 | -0.54 | 0.05 | 0.59 |
| JOI-E2 | SB-744941 | 0.65 | 0.27 | 0.46 | 0.19 |
| JOI-E3 | GW574782A | 2.94 | 2.33 | 2.635 | 0.305 |
| JOI-E4 | GW575808A | 0.72 | 0.75 | 0.735 | 0.015 |
| JOI-E5 | GW654652X | 0.57 | 0.21 | 0.39 | 0.18 |
| JOI-F1 | GW513184X | 2.98 | 2.77 | 2.875 | 0.105 |
| JOI-F2 | GW622055X | 0.96 | 0.35 | 0.655 | 0.305 |
| JOI-F3 | GW589961A | 4.23 | 2.87 | 3.55 | 0.68 |
| JOI-F4 | GW569293X | 0.64 | -0.06 | 0.29 | 0.35 |
| JOI-F5 | GSK466317A | 0.76 | 0.67 | 0.715 | 0.045 |
| JOI-G1 | GW682841X | 0.65 | 0.22 | 0.435 | 0.215 |
| JOI-G2 | GSK1713088A | 2.95 | 2.58 | 2.765 | 0.185 |
| JOI-G3 | GW819230X | 0.9 | 0.21 | 0.555 | 0.345 |
| JOI-G4 | GW769076X | 1.03 | 0.33 | 0.68 | 0.35 |
| JOI-G5 | GW276655X | 0.99 | 0.68 | 0.835 | 0.155 |
| JOI-H1 | GW770249A | 4.06 | 3.13 | 3.595 | 0.465 |
| JOI-H2 | SB-242721 | 1.08 | -0.14 | 0.47 | 0.61 |
| JOI-H3 | GW642138X | 3.15 | 2.45 | 2.8 | 0.35 |
| JOI-H4 | GW820759X | 0.76 | 0.5 | 0.63 | 0.13 |
| JOI-H5 | GW819077X | 1.3 | 1.52 | 1.41 | 0.11 |
| JOI-A6 | GW680191X | -0.44 | -0.29 | -0.365 | 0.075 |
| JOI-A7 | GW583373A | 1.12 | 1.14 | 1.13 | 0.01 |
| JOI-A8 | GW335962X | -0.12 | -0.73 | -0.425 | 0.305 |
| JOI-A9 | GW301784X | -0.7 | -0.01 | -0.355 | 0.345 |
| JOI-A10 | GW824645A | -0.05 | 0.41 | 0.18 | 0.23 |
| JOI-B6 | GSK554170A | -3.58 | -0.21 | -1.895 | 1.685 |
| JOI-B7 | GSK1220512A | 0.01 | 0.48 | 0.245 | 0.235 |
| JOI-B8 | SB-253226 | -0.19 | -0.65 | -0.42 | 0.23 |
| JOI-B9 | GW282536X | -0.12 | -0.09 | -0.105 | 0.015 |
| JOI-B10 | GI98581X | -0.07 | 0.53 | 0.23 | 0.3 |
| JOI-C6 | GSK579289A | -0.57 | -0.5 | -0.535 | 0.035 |
| JOI-C7 | GW779439X | 0.79 | 0.87 | 0.83 | 0.04 |
| JOI-C8 | GW734508X | 0.37 | -0.46 | -0.045 | 0.415 |
| JOI-C9 | GW852849X | -0.59 | -0.47 | -0.53 | 0.06 |
| JOI-C10 | GW683134A | 3.66 | 3.6 | 3.63 | 0.03 |
| JOI-D6 | GW576924A | 1.11 | 1.35 | 1.23 | 0.12 |
| JOI-D7 | GW771127A | 0.07 | -0.15 | -0.04 | 0.11 |
| JOI-D8 | SB-210313 | 0.51 | -0.6 | -0.045 | 0.555 |
| JOI-D9 | GW580496A | 2.21 | 2.11 | 2.16 | 0.05 |
| JOI-D10 | GSK2219385A | -1.72 | 0.37 | -0.675 | 1.045 |
| JOI-E6 | GW633459A | 0.75 | 0.38 | 0.565 | 0.185 |
| JOI-E7 | GW795493X | 2.14 | 3.26 | 2.7 | 0.56 |
| JOI-E8 | GSK614526A | -0.2 | -0.5 | -0.35 | 0.15 |
| JOI-E9 | GW568377B | -0.34 | -0.02 | -0.18 | 0.16 |
| JOI-E10 | GW305074X | 2.55 | 1.29 | 1.92 | 0.63 |
| JOI-F6 | GSK466314A | -0.16 | -0.4 | -0.28 | 0.12 |
| JOI-F7 | SB-250715 | -1.93 | -0.42 | -1.175 | 0.755 |
| JOI-F8 | GSK1007102B | -0.47 | -1.07 | -0.77 | 0.3 |
| JOI-F9 | GW693917X | 3.12 | 4.37 | 3.745 | 0.625 |
| JOI-F10 | GSK2110236A | 0.07 | 0.78 | 0.425 | 0.355 |
| JOI-G6 | GW615311X | 0.51 | 0.64 | 0.575 | 0.065 |
| JOI-G7 | SB-284847-BT | -0.18 | -0.15 | -0.165 | 0.015 |
| JOI-G8 | GSK2220400A | 0.26 | -0.29 | -0.015 | 0.275 |
| JOI-G9 | GSK635416A | -0.22 | 0.1 | -0.06 | 0.16 |
| JOI-G10 | SB-251527 | -0.04 | 0.47 | 0.215 | 0.255 |
| JOI-H6 | GW621823A | 0.69 | 0.67 | 0.68 | 0.01 |
| JOI-H7 | SB-254169 | -0.05 | 0.07 | 0.01 | 0.06 |
| JOI-H8 | SB-610251-B | 0.84 | 0.8 | 0.82 | 0.02 |
| JOI-H9 | SB-693162 | -0.26 | -0.11 | -0.185 | 0.075 |
| JOI-H10 | GW701427A | 0.55 | 1.14 | 0.845 | 0.295 |
| JOH-A1 | GW809885X | -0.07 | 0.72 | 0.325 | 0.395 |
| JOH-A2 | GW678313X | -0.31 | 0.32 | 0.005 | 0.315 |
| JOH-A3 | GW782912X | -0.99 | -0.18 | -0.585 | 0.405 |
| JOH-A4 | GW694234A | 2.52 | 3.93 | 3.225 | 0.705 |
| JOH-A5 | GW432441X | -0.41 | -0.35 | -0.38 | 0.03 |
| JOH-A6 | SB-251505 | 0.06 | -0.14 | -0.04 | 0.1 |
| JOH-A7 | GW683109X | -0.12 | -0.04 | -0.08 | 0.04 |
| JOH-A8 | SB-735467 | 1.06 | -0.36 | 0.35 | 0.71 |
| JOH-A9 | GW785404X | 0.87 | -0.46 | 0.205 | 0.665 |
| JOH-A10 | GW743024X | 0.61 | 0.15 | 0.38 | 0.23 |
| JOH-B1 | GW572401X | -0.88 | 0.23 | -0.325 | 0.555 |
| JOH-B2 | GW280670X | -0.55 | 0.17 | -0.19 | 0.36 |
| JOH-B3 | GW407323A | 1.1 | 0.26 | 0.68 | 0.42 |
| JOH-B4 | GW282449A | -0.87 | -0.24 | -0.555 | 0.315 |
| JOH-B5 | SB-347804 | -0.07 | 0.75 | 0.34 | 0.41 |
| JOH-B6 | SB-242718 | 0.12 | -0.15 | -0.015 | 0.135 |
| JOH-B7 | GW616030X | 1.59 | -0.44 | 0.575 | 1.015 |
| JOH-B8 | SB-358518 | -0.26 | -0.07 | -0.165 | 0.095 |
| JOH-B9 | SB-737198 | -0.37 | -0.41 | -0.39 | 0.02 |
| JOH-B10 | GSK994854A | 0.81 | 0.54 | 0.675 | 0.135 |
| JOH-C1 | GW406731X | -0.95 | -0.03 | -0.49 | 0.46 |
| JOH-C2 | GW279320X | -1.07 | 0.22 | -0.425 | 0.645 |
| JOH-C3 | SB-698596-AC | -0.59 | 0.16 | -0.215 | 0.375 |
| JOH-C4 | GW794607X | -0.78 | 0.42 | -0.18 | 0.6 |
| JOH-C5 | SB-476429-A | -1.64 | -0.86 | -1.25 | 0.39 |
| JOH-C6 | GW817394X | -0.34 | 0.85 | 0.255 | 0.595 |
| JOH-C7 | GW693881A | 4.7 | 4.61 | 4.655 | 0.045 |
| JOH-C8 | SB-772077-B | -0.58 | -0.23 | -0.405 | 0.175 |
| JOH-C9 | SB-431542-A | 0.16 | 0.51 | 0.335 | 0.175 |
| JOH-C10 | GSK238063A | 1.22 | 1.63 | 1.425 | 0.205 |
| JOH-D1 | GW780056X | -0.86 | -0.01 | -0.435 | 0.425 |
| JOH-D2 | GW811168X | -0.46 | 1.35 | 0.445 | 0.905 |
| JOH-D3 | GW659893X | 3.96 | 3.72 | 3.84 | 0.12 |
| JOH-D4 | GW829874X | -0.62 | 0.51 | -0.055 | 0.565 |
| JOH-D5 | GSK270822A | -0.77 | 0.16 | -0.305 | 0.465 |
| JOH-D6 | GW396574X | 1.36 | 1.57 | 1.465 | 0.105 |
| JOH-D7 | SB-736302 | -0.37 | -0.17 | -0.27 | 0.1 |
| JOH-D8 | GSK269962B | 0.38 | 0.87 | 0.625 | 0.245 |
| JOH-D9 | GW632580X | 0 | 0.13 | 0.065 | 0.065 |
| JOH-D10 | GSK312948A | 0.12 | -0.11 | 0.005 | 0.115 |
| JOH-E1 | GSK248233A | -0.09 | -0.5 | -0.295 | 0.205 |
| JOH-E2 | GW442130X | -0.35 | 0.48 | 0.065 | 0.415 |
| JOH-E3 | GW693481X | -1.1 | 0.25 | -0.425 | 0.675 |
| JOH-E4 | GW679410X | -1.21 | 0.43 | -0.39 | 0.82 |
| JOH-E5 | GW805758X | -0.31 | 0.99 | 0.34 | 0.65 |
| JOH-E6 | GW458787A | 0.84 | 0.98 | 0.91 | 0.07 |
| JOH-E7 | GW830900A | -0.27 | 0.41 | 0.07 | 0.34 |
| JOH-E8 | GW445014X | -1.29 | -0.41 | -0.85 | 0.44 |
| JOH-E9 | SB-725317 | 0.05 | 0.43 | 0.24 | 0.19 |
| JOH-E10 | GW459057A | 0.4 | 0.2 | 0.3 | 0.1 |
| JOH-F1 | GW683768X | -0.43 | 0.23 | -0.1 | 0.33 |
| JOH-F2 | GW275616X | -0.88 | 0.23 | -0.325 | 0.555 |
| JOH-F3 | GW642125X | -0.41 | 0.57 | 0.08 | 0.49 |
| JOH-F4 | GW300660X | -0.66 | 0.67 | 0.005 | 0.665 |
| JOH-F5 | GW549034X | -1.03 | 0.09 | -0.47 | 0.56 |
| JOH-F6 | GW708336X | -0.66 | 0.28 | -0.19 | 0.47 |
| JOH-F7 | GW782907X | -1.07 | 0.7 | -0.185 | 0.885 |
| JOH-F8 | SB-743899 | -0.69 | -0.23 | -0.46 | 0.23 |
| JOH-F9 | GSK980961A | -0.74 | 0.22 | -0.26 | 0.48 |
| JOH-F10 | SB-590885-R | 0.15 | 0.02 | 0.085 | 0.065 |
| JOH-G1 | GSK949675A | -1.61 | -1.11 | -1.36 | 0.25 |
| JOH-G2 | GW786460X | -1.42 | 0.43 | -0.495 | 0.925 |
| JOH-G3 | GW876790X | -1.06 | 0.49 | -0.285 | 0.775 |
| JOH-G4 | GW427984X | -0.74 | 0.3 | -0.22 | 0.52 |
| JOH-G5 | GSK2186269A | -1.79 | -1.53 | -1.66 | 0.13 |
| JOH-G6 | GW684626B | -0.58 | 0.79 | 0.105 | 0.685 |
| JOH-G7 | GW806290X | -0.25 | -0.28 | -0.265 | 0.015 |
| JOH-G8 | GW445012X | -1.94 | -0.02 | -0.98 | 0.96 |
| JOH-G9 | GW711782X | -0.32 | 0.47 | 0.075 | 0.395 |
| JOH-G10 | GW576609A | 0.92 | 1.03 | 0.975 | 0.055 |
| JOH-H1 | GSK1819799A | 0.28 | 0.78 | 0.53 | 0.25 |
| JOH-H2 | GSK625137A | -1.09 | 0.28 | -0.405 | 0.685 |
| JOH-H3 | GW801372X | -1.25 | 0.66 | -0.295 | 0.955 |
| JOH-H4 | GSK317314A | -2.1 | 0.14 | -0.98 | 1.12 |
| JOH-H5 | GW284372X | 0.38 | 0.28 | 0.33 | 0.05 |
| JOH-H6 | GW701032X | -1.34 | 0.38 | -0.48 | 0.86 |
| JOH-H7 | SB-216385 | -0.55 | 0.47 | -0.04 | 0.51 |
| JOH-H8 | GW439255X | -0.51 | 0.95 | 0.22 | 0.73 |
| JOH-H9 | GW618013X | 0.44 | 0.76 | 0.6 | 0.16 |
| JOH-H10 | GSK1326255A | 2.16 | 2.18 | 2.17 | 0.01 |
| TRB2+ATP+EDTA |  | 2.325 | 2.78 | 2.5525 | 0.2275 |

